# Biophysical trade-offs in antibody evolution are resolved by conformation-mediated epistasis

**DOI:** 10.64898/2026.03.12.711465

**Authors:** Cole R. Tharp, Claudio Catalano, Anthony Khalifeh, Sam Ghaffari-Kashani, Ruimin Huang, Gyunghoon Kang, Giovanna Scapin, Angela M. Phillips

## Abstract

Protein evolution is constrained by multidimensional biophysical factors, in which mutations that enhance one property often compromise another. Antibodies represent an extreme case: they evolve rapidly to bind diverse antigens, yet mutations that improve affinity can disrupt folding, reduce cell-surface trafficking, or promote self-reactivity, and are typically selected against during affinity maturation. Though biophysical characterization of individual antibodies suggests that such trade-offs are pervasive, their impact on antibody evolutionary trajectories remains unclear, in part because existing high-throughput biophysical methods rely on heterologous systems that are often poorly suited for human proteins. Here, we develop a high-throughput platform to quantify multiple biophysical parameters of large libraries of full-length proteins that are natively synthesized, processed, and displayed on human cells. We apply this approach to a human antibody lineage that matures to recognize divergent SARS-CoV-2 variants by measuring the surface expression, antigen affinity, and self-reactivity for all 2^13^ possible evolutionary intermediates between the unmutated and mature sequences. These measurements reveal that mutations differentially affect these biophysical properties – in some cases, improving one property at the expense of another. We leverage these data to compute the likelihood of all possible evolutionary paths, finding that very few paths can navigate these multidimensional requirements. The few accessible paths acquire mutations in a specific order that either circumvent trade-offs between biophysical properties or offset deleterious effects on one property with beneficial effects on another. By determining the structures of the ancestral and evolved antibodies, we find that these coordinated mutational effects arise from a conformational rearrangement that alleviates steric clashes and reshapes the biophysical landscape, enabling otherwise inaccessible mutational paths. Together, this work defines the multidimensional biophysical constraints and structural mechanisms that govern antibody evolution and establishes a general framework for mapping and predicting the biophysical effects of mutations in human proteins.

## Introduction

Mutations rarely impact a single property of a protein – instead, they alter many distinct biophysical properties that jointly determine evolutionary outcomes^1–4^. Across diverse systems, mutations can have differential, or pleiotropic, effects on protein stability, folding, binding specificity, and catalysis, resulting in biophysical trade-offs that restrict the accessibility of adaptive mutational trajectories^1–3,5–7^. These effects may be especially consequential for proteins with multiple functions or stringent quality-control requirements, particularly when evolving traits that require many mutations. This challenge raises the question we focus on here: how do proteins navigate multidimensional constraints to evolve new functions?

Antibodies represent one such class of proteins, rapidly acquiring mutations that have pleiotropic biophysical effects^7,8^. During affinity maturation, B-cell receptors (membrane-bound antibodies) acquire somatic mutations and undergo selection to bind antigens, resulting in antibody sequences with improved antigen affinities^9–11^. Because B-cell proliferation is tightly coupled to the number of antigen molecules bound, it is, in principle, determined by the number of B-cell receptors, their affinity to antigen, and competitive interactions with non-cognate antigens^7,9,11,12^. Thus, the differential effects of mutations on antibody surface expression, binding affinity, and specificity likely impact antibody mutational trajectories. These multi-dimensional constraints may help explain why broadly neutralizing antibodies (bnAbs), which acquire many mutations to bind divergent pathogens^13–16^, arise at low frequencies in human repertoires^17–20^. Despite the potential importance of biophysical trade-offs in shaping immunity and informing vaccine design, we do not understand the extent to which mutations alter antibody biophysics and how such effects limit or potentiate antibody maturation.

Although few studies have dissected the biophysical effects of mutations in antibodies, they show that mutations can differentially impact distinct biophysical properties – for example, improving antigen affinity at the cost of antibody expression. In a dataset of ∼400 human B-cell-derived antibodies, Shehata et al found that somatically mutated antibodies tend to exhibit reduced thermal stability compared to naïve antibodies^8^. Though stability generally declined with increasing mutational load, all antibodies they examined exceeded a stability threshold, suggesting that a minimum level of stability may be required for antibody folding. Consistent with this possibility, other studies of individual antibody lineages demonstrate that affinity-enhancing mutations can be destabilizing and require compensatory mutations to maintain sufficient stability^21^. Importantly, however, B-cell proliferation is influenced by the number of B-cell receptors on the cell surface^9,12^, which is determined not only by antibody stability but also by cellular folding and trafficking^8,22,23^. The few studies that have characterized B-cell receptor surface expression have observed that highly expressed receptors tend to have fewer somatic mutations^8^, implying that affinity-enhancing mutations compromise folding or trafficking. Together, existing datasets indicate that mutational effects on surface expression (via stability, folding, or trafficking) impose stringent biophysical constraints. Still, their limited scope and resolution leave open fundamental questions about how these properties shape the accessibility of antibody mutational trajectories.

Somatic mutations can also affect antibody self-reactivity, encompassing both specific (*i.e.,* autoreactive) and nonspecific (*i.e.,* polyspecific) interactions with self-molecules. These effects have been observed in directed evolution campaigns^24^, longitudinal studies in mice^7,25^, and human B-cell repertoires^8,26–28^, and, further, are often selected against during *in vivo* affinity maturation^7,25,29^. Conversely, moderate levels of self-reactivity may be beneficial. For example, polyspecificity can correspond to increased flexibility, enabling antibodies to recognize sites that are sterically occluded or rapidly evolving^27,28,30^. Thus, although self-reactivity is readily altered by mutation and subject to selection, the frequency with which these mutations arise and their influence on antibody evolutionary trajectories remain unresolved.

A major barrier to addressing these questions is the lack of high-throughput biophysical methods for assaying full-length antibodies in their native IgG format. Existing platforms, such as yeast and phage display, enable high-throughput binding measurements but rely on truncated antibody constructs that undergo distinct post-translational modifications and processing compared with full-length antibodies synthesized in human cells^31–33^. These differences can confound measurements of antigen affinity and specificity, as truncated antibody constructs can exhibit divergent binding behavior^34,35^. Furthermore, because antibody surface expression depends on antibody stability, folding, and trafficking, it is strongly influenced by cellular protein-folding and quality-control factors that vary across heterologous systems^36–39^. Collectively, these differences are known to alter antibody conformation and biophysical properties, limiting the application of these methods for assessing how mutations affect full-length IgGs.

To overcome these limitations, we developed BioPhy-Seq, a high-throughput platform that enables quantitative measurement of multiple biophysical parameters for large libraries of full-length IgG antibodies synthesized and processed in human cells. This approach makes it possible, for the first time, to systematically examine how mutations affect surface expression, affinity, and self-reactivity across antibody evolutionary trajectories. Further, we note that this method can be applied to measure any fluorescence-coupled property for any human membrane protein. Compared to existing human cell-display methods^40–43^, BioPhy-Seq uses titrations to measure absolute biophysical properties (*e.g.,* equilibrium binding affinities, *K_D_*, and half-maximal effective concentrations, *EC_50_*), enabling comparisons across datasets that are challenging with relative measurements. Here, we apply BioPhy-Seq to interrogate an antibody lineage that acquires breadth to divergent SARS-CoV-2 variants, generating all 2^13^ possible evolutionary intermediates between the germline (*i.e.,* unmutated) and mature sequences. For this combinatorial library, we measure affinity to three divergent SARS-CoV-2 spike variants, surface expression, and polyspecificity (this lineage is not known to exhibit autoreactivity)^44,45^. By pairing these comprehensive biophysical data with structural analyses, we define how biophysical properties – and trade-offs between them – shape the accessibility of antibody mutational trajectories and establish a broader framework for understanding and predicting the biophysical effects of mutations in human proteins.

## Results

### A multidimensional biophysical landscape reveals trade-offs in an antibody lineage

To examine whether antibody mutational pathways vary in their biophysical accessibility (**Fig. 1a**), we comprehensively profiled all 2^13^ possible evolutionary intermediates of the Omi32 antibody lineage, which matured to recognize divergent variants of the SARS-CoV-2 spike receptor binding domain (RBD) by acquiring 13 mutations^44^. To achieve this scale, we developed BioPhy-Seq, a high-throughput platform that measures biophysical properties across large libraries of full-length human proteins. Briefly, this method leverages genomic landing pads^42,46^ to generate libraries of human cells, in which each cell expresses a single antibody sequence on its surface (**ED Fig. 1**). Cell libraries are subjected to fluorescence-coupled treatments that vary depending on the desired biophysical property (*e.g.,* antigen affinity is measured by titrating fluorescently labeled antigen). Biophysical measurements are then made in bulk by combining fluorescence-activated cell sorting with deep sequencing^33,47^.

**Figure 1.**
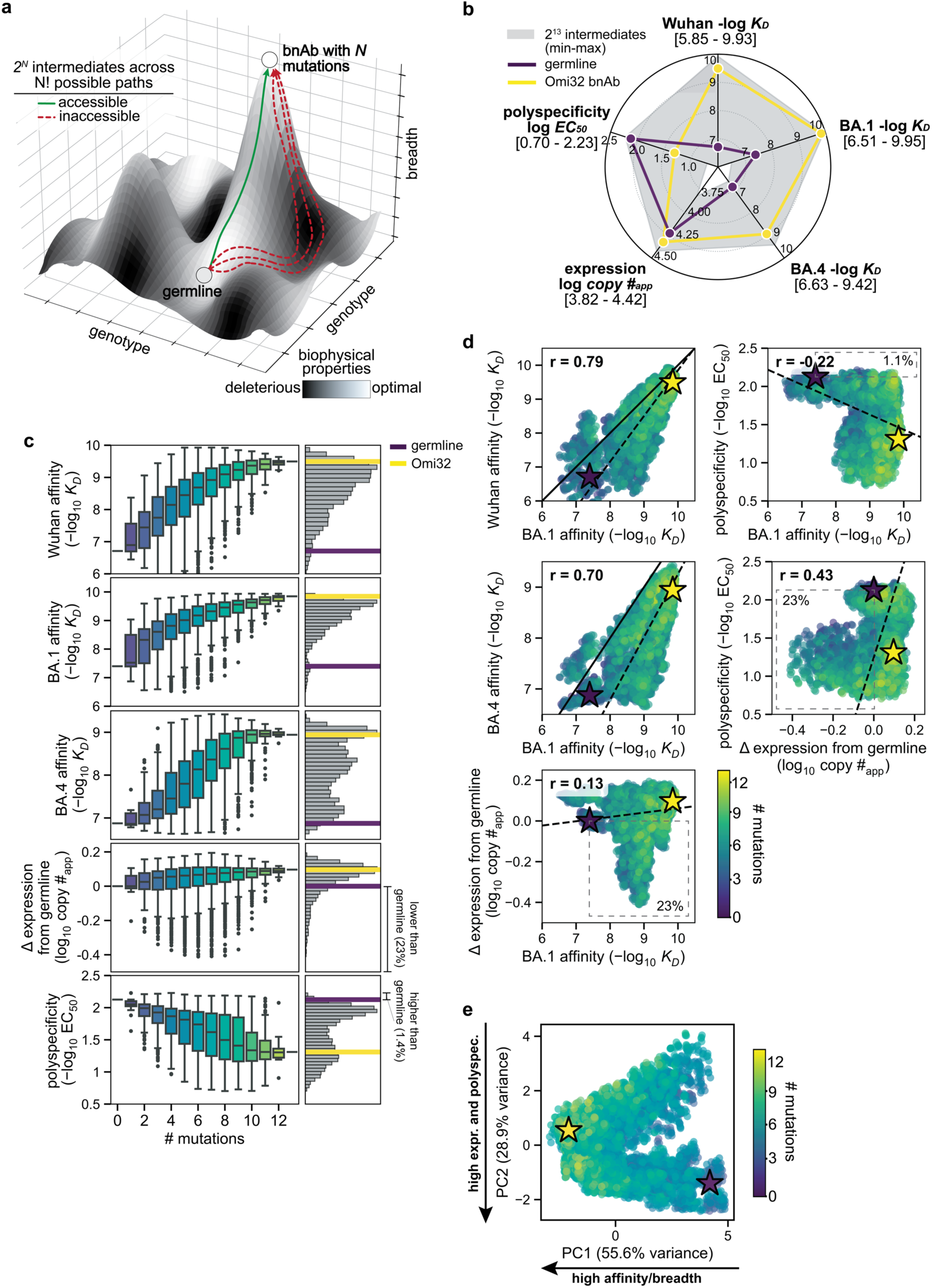
| Multidimensional biophysical constraints shape the evolutionary landscape of a human antibody lineage. **a,** Conceptual schematic illustrating how mutations can alter biophysical properties that collectively constrain the accessibility of mutational paths to bnAbs. Mutations that improve breadth may compromise another biophysical property, generating trade-offs that restrict viable evolutionary trajectories. **b,** Overview of the combinatorially complete library comprising all 2¹³ intermediates between the germline and mature states of the Omi32 antibody lineage. Five biophysical properties (affinity to Wuhan, BA1, BA4, surface expression, and polyspecificity) were measured for all 2^13^ intermediates. The range across the library is shown in gray, with the germline (purple) and Omi32 (yellow) sequences annotated. **c,** Left: Box plots of measured biophysical properties show variation with respect to mutagenic load. Right: Histograms show variation across the library, with germline (purple) and Omi32 (yellow) sequences annotated. **d,** Pairwise relationships between biophysical properties show correlations and trade-offs, including correlations between affinity to different antigens and trade-offs between polyspecificity and expression. Omi32 and germline are labeled as yellow and purple stars, respectively. Pearson’s correlation coefficient (r), solid identity line (1:1), and dashed least-squares regression line are shown on plot. **e,** Low-dimensional embedding (via principal component analysis) of the five-dimensional landscape highlights variation structured by two principal components, one corresponding to antigen affinity (and hence, breadth, PC1: 55.6% variance), and another corresponding to expression and polyspecificity (PC2: 28.9% variance). **To include all 8,192 genotypes, phenotypes predicted from linear models were used for analysis and plotting in b-e (see Methods and comparison to experimental phenotypes (ED Fig. 1)).

Here, BioPhy-Seq empowered comprehensive measurement of antibody surface expression, affinity to divergent SARS-CoV-2 RBD variants, and polyspecificity (**ED Fig. 1**). Across all properties, evolutionary intermediates spanned a broad dynamic range (**Fig. 1b**), with many intermediates exhibiting impaired biophysical properties compared to the germline, revealing that antibody maturation is shaped by multidimensional biophysical constraints that are challenging to measure reliably in heterologous display systems.

For affinity to Wuhan, BA1, and BA4 RBD, most intermediates exhibited affinities between the germline precursor and Omi32, with a small fraction exceeding the affinity of the mature antibody (**Fig. 1b-c**). Furthermore, across all antigens, affinity increased gradually with the number of mutations. For Wuhan and BA1, even those with relatively few mutations exhibited comparable affinity to Omi32, suggesting that early mutational steps can substantially improve affinity. In contrast, affinity to BA4, which carries the L452R mutation in the epitope recognized by Omi32^48^, generally required more mutations.

Unlike the multimodal affinity distributions, expression exhibited a Poisson-like distribution. Though germline and Omi32 have similar expression levels, a substantial portion of the library (23%) has reduced expression compared to germline. Interestingly, expression is not simply correlated with the number of mutations, as poorly expressed variants contain varying numbers of mutations. We find that BioPhy-Seq measurements of expression on HEK cells reflect measurements made on B-cells, and further, that surface expression is an aggregate measure of antibody stability, folding, and trafficking, and thus is not a simple proxy for thermal stability (**ED Fig. 2**)^23^.

**Figure 2.**
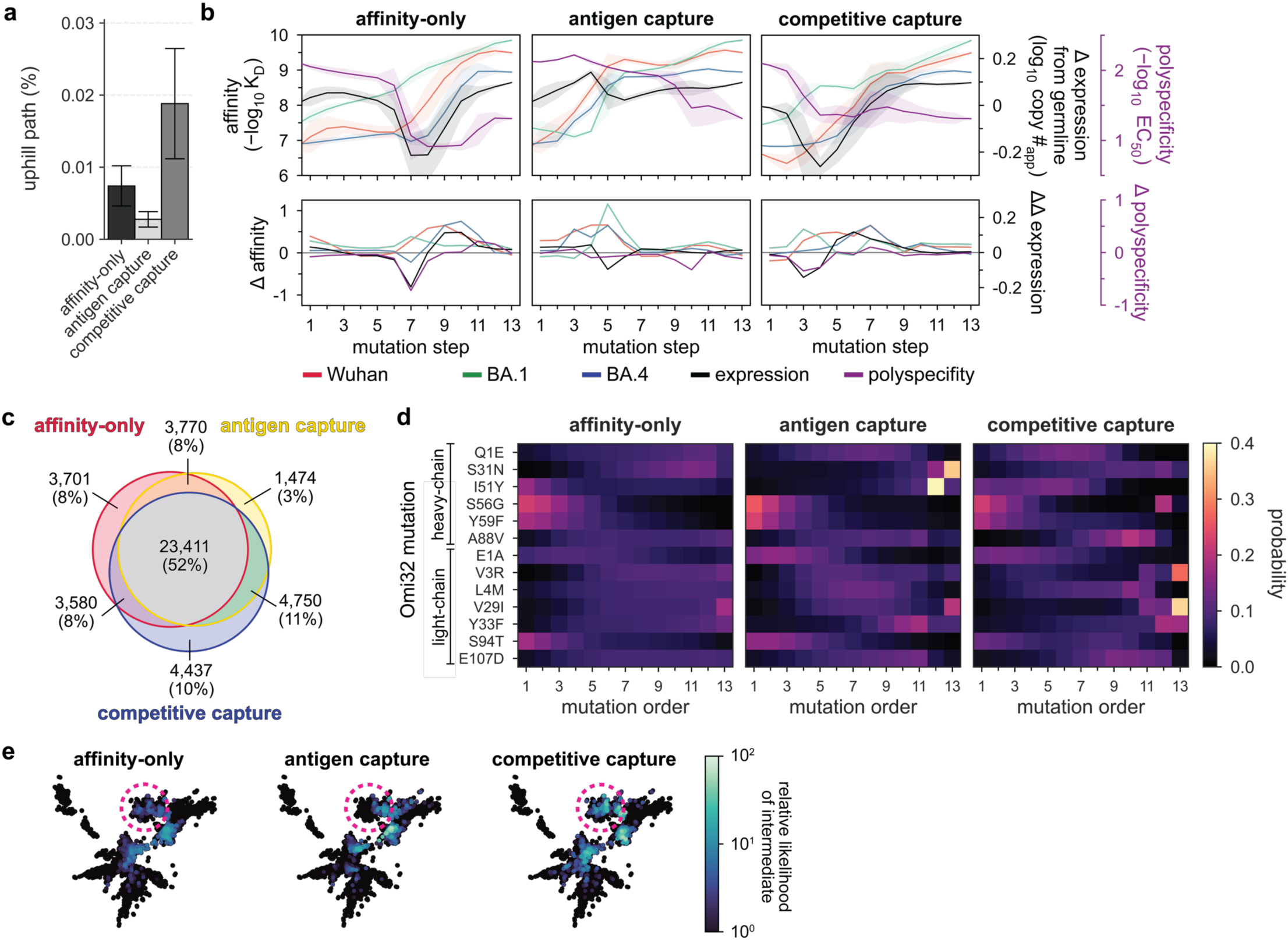
| Biophysical trade-offs constrain evolutionary path accessibility. **a,** Percentage of all possible paths (13!) that are uphill (*i.e.,* where fitness improves at every mutational step) under distinct selection models. Affinity-only: selection is solely based on improvements in affinity, antigen capture: selection is determined equally by changes in affinity and expression, competitive capture: selection is determined equally by changes in affinity, expression, and polyspecificity (see Methods for details). **b,** Top: mean phenotype at each mutational step for the most likely 1,000 paths under the three selection models (here, all models implement weak selection pressure where neutral and deleterious mutations are disfavored but allowed – see Methods for details). The shaded area represents one standard deviation from the mean. Bottom: mean change in phenotype at each step. **c,** Venn diagram of mutations in uphill paths shared across the three models. Total number and percentage of uphill mutational steps is annotated. **d,** Normalized likelihood that each mutation occurred at a particular mutational step for each of the weak selection models. **e,** Likelihood of each intermediate is projected onto a force-directed graph, where each antibody sequence is represented by a node, single-mutant neighbors are connected by edges (not shown) that are weighted by the difference in BA1 affinity. Clustering is driven by a force that is inversely proportional to the change in BA1 affinity, such that sequences with more similar affinities cluster together. Distinct clusters represent antibody sequences with distinct sets of mutations. One example of a cluster (*i.e.,* sequences that share a specific set of mutations that define their affinity to BA1) that differs in likelihood across the models is annotated (pink dotted line). **To include all 8,192 genotypes, phenotypes predicted from linear models were used for analysis and plotting in a-e (see Methods and comparison to experimental phenotypes (ED Fig. 1)).

In contrast to expression, polyspecificity exhibits a multimodal distribution, with germline amongst the most polyspecific sequences in the library, and Omi32 exhibiting substantially reduced polyspecificity. Furthermore, polyspecificity gradually improves (declines) with increasing mutation number, although a small subset of sequences (1.4%) exhibits elevated polyspecificity, indicating that certain combinations of mutations may be disfavored by selection.

Correlations across biophysical properties revealed distinct relationships (**Fig. 1d**). Affinities to Wuhan, BA1, and BA4 were positively correlated, though affinity to Wuhan and BA1 exhibited higher correlation than BA1 and BA4, likely reflecting the L452R mutation in the BA4 epitope. In contrast, we observe much weaker relationships between affinity and expression, where, although expression tends to rise with affinity, there are many (23%) variants where improved affinity accompanies reduced expression, suggesting a trade-off between improving affinity and maintaining proper antibody folding and trafficking. Generally, polyspecificity improves with affinity, with exceedingly few variants (1.1%) having improved affinity at the expense of polyspecificity. Further, we find that many variants with improved polyspecificity have reduced expression (23%). Together, these patterns reveal a highly structured biophysical landscape in which the evolution of affinity and breadth is coupled to an improvement in polyspecificity and, at times, a reduction in expression.

To integrate variation across all measured properties, we performed a principal component analysis of the biophysical data (**Fig. 1e**)^49^. This analysis revealed that variation across the library is dominated by two principal axes corresponding to *(1)* affinity to divergent antigens (breadth), and *(2)* an integrated expression-polyspecificity dimension. Intermediates clustered into two populations, one corresponding to higher polyspecificity and expression than the other, with each population spanning a continuum of breadth. Along this trajectory, the degree of somatic mutation was associated with increased breadth, with highly mutated variants exhibiting a narrower range of polyspecificity and expression than less mutated variants. Collectively, these results show that the maturation of this antibody lineage is jointly shaped by affinity, expression, and polyspecificity, motivating the following analysis of how these pleiotropic effects shape evolutionary trajectories.

### Biophysical constraints shape evolutionary paths to antibody breadth

The quantitative nature of our combinatorial dataset enables direct evaluation of the biophysical accessibility of all possible 13! (6e9) mutational trajectories from the germline precursor to Omi32. To this end, we applied simple population-genetic models of selection^50,51^– intended not to recapitulate the full complexity of selection in germinal centers (*e.g.,* T-cell help, spatial structure, *etc.*)^9,52–54^, but rather to define how biophysical properties alone can shape the accessibility of antibody evolutionary paths. Unlike prior analyses based on yeast display, which only considered mutational effects on antigen affinity^55,56^, our BioPhy-Seq datasets allow us to incorporate additional constraints arising from antibody expression and polyspecificity, both of which are known to influence selection during affinity maturation^7,12,28^. Because B-cell proliferation is tightly coupled to the total number of antigen molecules bound, it is dependent on antibody surface expression, antigen affinity, and non-specific interactions that may block antigen binding^7,9,12^. To dissect the contributions of these biophysical properties, we consider three models of varying complexity.

Each model considers mutational effects on one or more biophysical parameters to compute the likelihood of all possible paths from germline to Omi32 (see Methods for details). First, we consider an affinity-only model, in which the probability of a mutation depends solely on its effect on affinity. Second, we construct an antigen-capture model in which mutational probabilities are defined by their collective impact on affinity and surface expression (*i.e.,* deleterious effects on affinity can be compensated for by gains in expression, and vice versa). Finally, we incorporate our polyspecificity data to construct a competitive-capture model, in which non-specific binding events can ‘compete’ with antigen binding, and mutational probabilities are thus equally contingent on affinity, expression, and polyspecificity. For each of these models, we consider selection with BA1, as this antigen likely drove the maturation of the Omi32 lineage, which was isolated following a BA1 breakthrough infection^44,45^. Additionally, we consider varying levels of selection stringency, ranging from strong selection, where all properties must improve at each mutational step, to moderate and weak selection, where neutral and even slightly deleterious mutations are tolerated^50,51^.

These analyses reveal that regardless of the selection model, pathways to the Omi32 sequence are highly constrained (**Fig. 2a**). For example, in the affinity-only model, exceedingly few paths are uphill (0.007%, ∼4.6e5 paths); this number declines slightly for the antigen-capture model (0.003%, ∼1.7e5 paths) and, intriguingly, increases for the competitive-capture model (0.02%, ∼1.2e6 paths). These differences are reflected in the most likely paths for each model (**Fig. 2b**), as well as the shared mutations across uphill paths (**Fig. 2c**). For the affinity-only model, the most likely paths consistently gain affinity to BA1 (and correspondingly, to Wuhan and BA4) while improving in polyspecificity and navigating an expression valley (**Fig. 2b**). In contrast, in the antigen-capture model, which incorporates effects on affinity and expression, the most likely paths maintain expression, coupling reductions in expression with gains in affinity and vice versa (**Fig. 2b**), resulting in fewer total uphill paths (**Fig. 2a,c**). In the competitive-capture model, improvement in polyspecificity can compensate for reductions in expression and affinity – including an expression valley that is absent in the most likely paths from the antigen-capture model (**Fig. 2b**) – increasing the total number of uphill paths (**Fig. 2a,c**). Notably, for each model, the least likely paths are also phenotypically distinct (**ED Fig. 3a**). In the affinity-only model, the least likely paths do not cross an expression valley, though they also exhibit less consistent improvements in affinity, reducing their overall likelihood. In the antigen-capture and competitive-capture models, the least likely paths improve in affinity quite early. In the former model, this gain is negated by costs to expression, and in the latter, by the lack of phenotypic improvement from subsequent mutagenesis. These differences between the most and least likely paths reflect coupled changes across properties. For example, mutational effects on expression and affinity are anticorrelated for the most likely paths, particularly in the antigen-capture model, and less so for the least likely paths, demonstrating that consistent gains in affinity are coupled to reductions in expression (**ED Fig. 3b**). Together, this analysis reveals that the accessibility of mutational trajectories is defined by coordinated changes across multiple properties.

**Figure 3.**
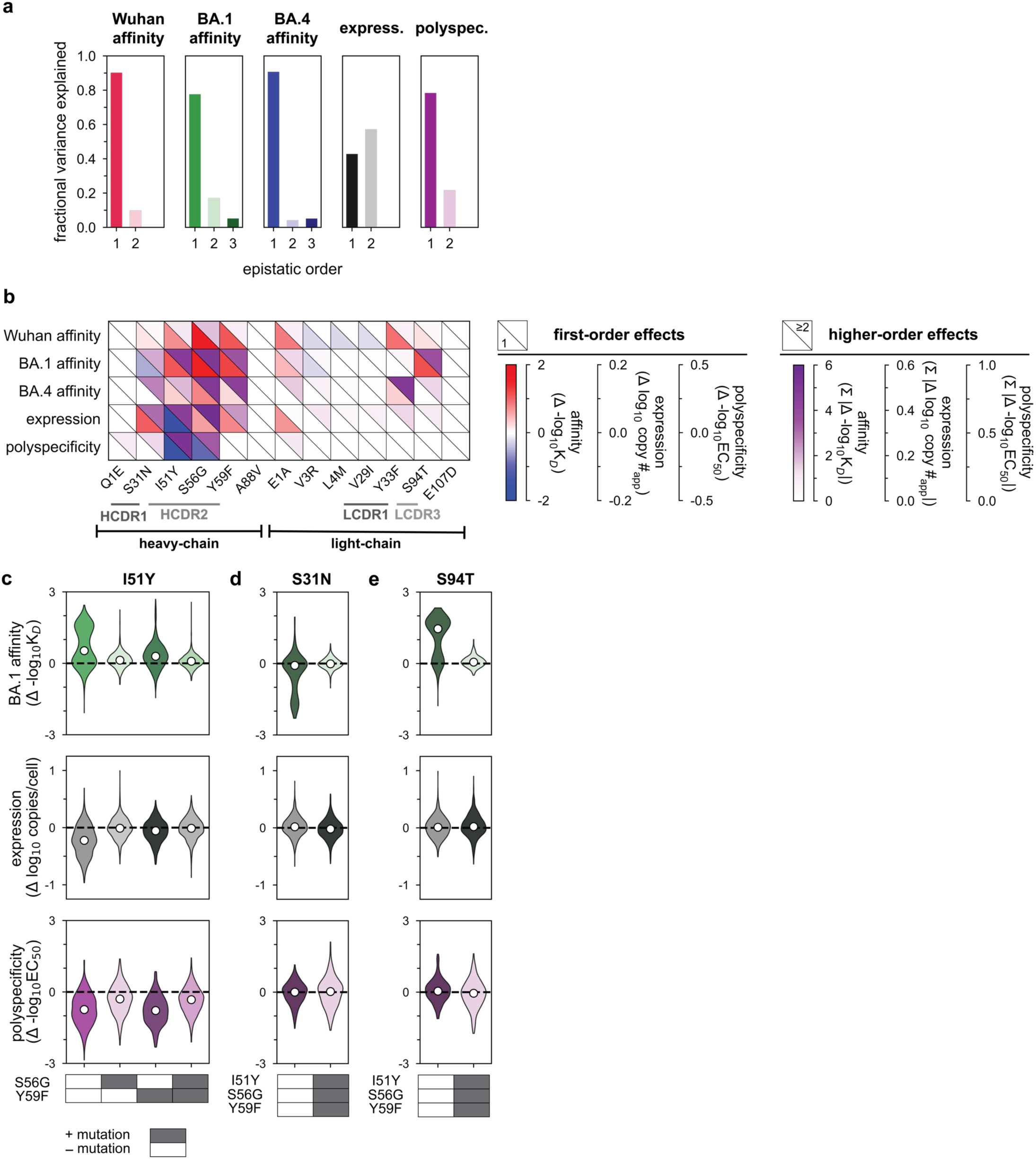
| Epistasis is significant for each biophysical property. **a,** Decomposition of variance for each biophysical property into additive and epistatic contributions for the optimal order model (using the reference-free framework, see Methods). **b,** Additive and epistatic effects (reference-based) of each Omi32 mutation on each biophysical property. The left triangle shows significant additive effects (Bonferroni-corrected *p*-value); the right triangle shows the sum of the absolute value of Bonferroni-significant epistatic coefficients (≥ order 2). All coefficients are presented in Extended Data 5-6 – these models were used to predict phenotypes for all 8,192 genotypes, which were used for the analysis and plotting in Figures 1 and 2. HCDR1/2: heavy-chain complementarity determining region 1/2. LCDR1/3: light-chain complementarity determining regions 1/3. **c,** Effect of I51Y on experimental measurements of BA1. affinity, expression, and polyspecificity for sequences with and without S56G and Y59F (N∼1024 genotypes per violin). **d,** Effect of S31N on experimental measurements of BA1. affinity, expression, and polyspecificity for sequences with and without I51Y, S56G, and Y59F (N∼512 genotypes per violin). **e,** Effect of S94T on experimental measurements of BA1. affinity, expression, and polyspecificity for sequences with and without and I51Y, S56G, Y59F (N∼512 genotypes per violin). **In c-e, the black line indicates zero effect. White dots indicate the median for each violin. Boxes underneath the plots indicate the presence (gray) or absence (white) of the specified mutation.

When we examine the likelihood of specific mutations along these trajectories, we observe differences across models. Though many mutations behave similarly across the models – with some preferred early (*e.g.,* S56G, Y59F, S94T) and others later on (e.g., S31N and most of the light chain mutations) – I51Y dramatically shifts from an early mutation in the affinity-only model, to a late mutation in the antigen-capture model, to an intermediate mutation in the competitive-capture model (**Fig. 2d**). This strong preference in mutational order shows how mutational effects are interdependent, or epistatic, and the shift in preference across the models demonstrates that mutations like I51Y differentially impact distinct biophysical properties, or are pleiotropic. Further, upon examining the likelihood of all possible 2^13^ intermediates, we find that those with high likelihood fall within small clusters of genetically similar sequences (**Fig. 2e**). Although many of these clusters overlap across models, others do not, again suggesting that mutational effects are both epistatic and pleiotropic.

### Epistasis mediates accessibility of biophysically-constrained paths

To examine the molecular basis for the preferred order of mutations, we used linear models to formally disentangle the effects of individual mutations and their combinations^57^. Broadly, mutational effects on all measured biophysical properties were highly background-dependent, revealing pervasive non-additivities (*i.e.,* epistasis). Epistasis was especially important for BA1 affinity, explaining 22% of the variation in affinity (**Fig. 3a**). For Wuhan and BA4, epistasis was less important, explaining 10% and 9% of the variation in affinity, respectively. For expression and polyspecificity, linear models had notably worse performance (**ED Fig. 4a**)– for expression, this is likely due to low variation across the library; for polyspecificity, this is likely due to higher experimental noise associated with these measurements (**ED Fig. 1**). Still, we find that both expression and polyspecificity are best described by a second-order model, with pairwise coefficients explaining 57% and 22% of the variance, respectively (**Fig. 3a**).

**Figure 4.**
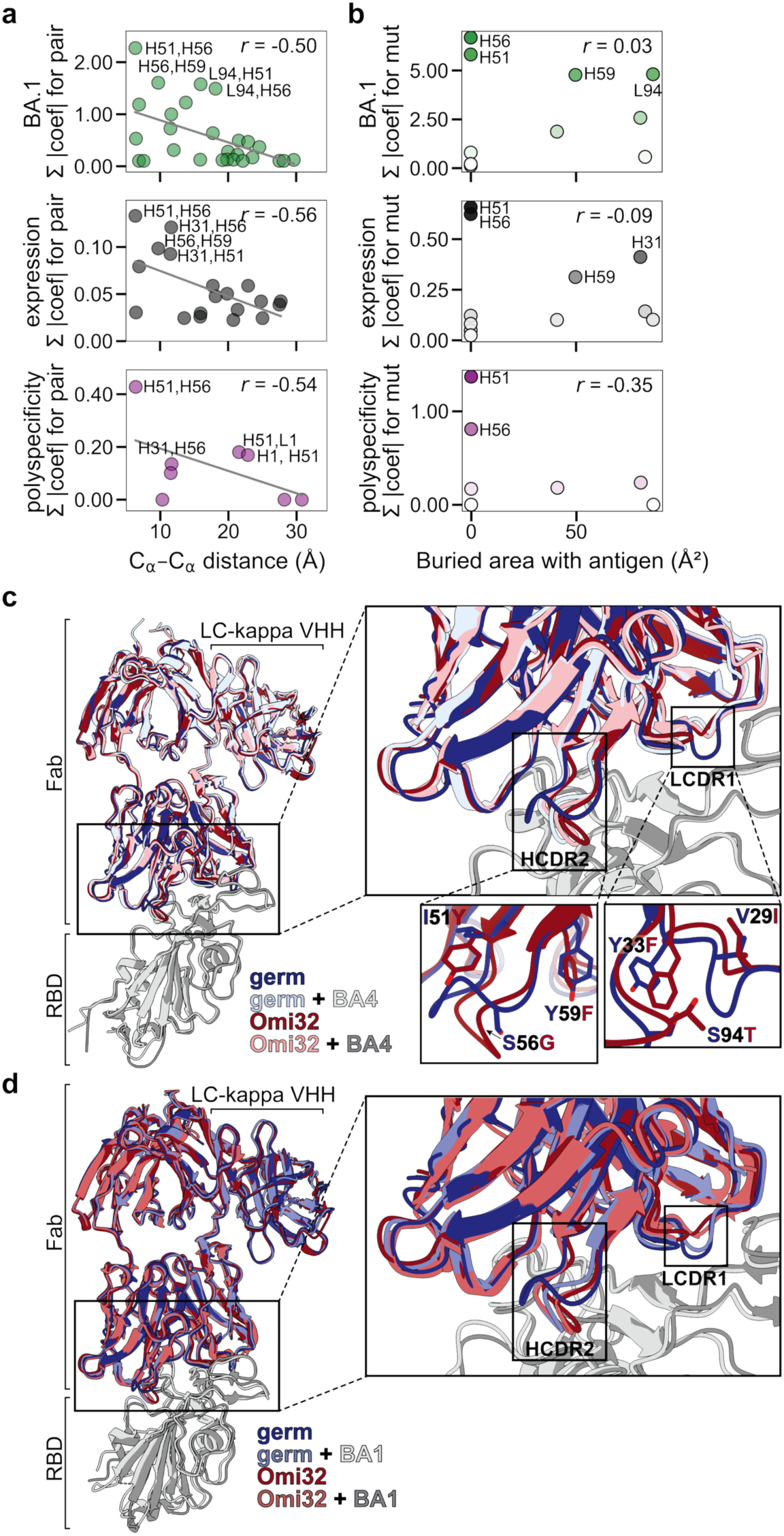
| Conformational rearrangements resolve steric conflicts and explain epistasis and trade-offs. **a,** Relationship between epistatic effects and distance between pairs of residues, computed from Omi32-BA1 structure (PDB ID 7ZFE)^44^. Strong interactions are labeled by chain (H: heavy; L: light) and residue number. Pearson r annotated on plot. **b,** Relationship between mutational effect size and buried area with the antigen, computed from Omi32-BA1 structure (PDB ID 7ZFE)^44^. Large-effect mutations are labeled by chain (H: heavy; L: light) and residue number. All mutations are shaded by total effect size. Pearson r annotated on plot. **c,** Cryo-EM structures of germline and Omi32, alone and in complex with BA4. Overlay highlights HCDR2 and LCDR1 loop rearrangements that accommodate mutated residues, resolving potential clashes. RMSD relative to Omi32: 0.866 Å (germline), 0.792 Å (germline+BA4), 0.612 Å (Omi32+BA4) for global alignment of light and heavy chains. **d**, Cryo-EM structures of germline and Omi32, alone and in complex with BA1. RMSD relative to Omi32: 0.866 Å (germline), 0.716 Å (germline+BA1), 0.791 Å (Omi32+BA1, 7ZFE) for global alignment of light and heavy chains. Structures for germline, Omi32, germline+BA4, Omi32+BA4, and germline+BA1 were determined to 3.1 Å, 3.2 Å, 3.4 Å, 3.2 Å, 3.2 Å nominal resolution, respectively. For loop rearrangements shown in c-d, other loop conformations were not observed during processing, see **ED Fig. 9-10**.

Across all five properties, we find that mutations with the largest effects reside within (or near) the complementarity-determining region (CDR) loops of the heavy and light chains. Generally, these mutations exhibit strong linear and epistatic effects (**Fig. 3b**, **ED Fig. 5-6**), demonstrating that while mutations can individually impact these properties, their combined effects are interdependent. Intriguingly, these same mutations exhibit strong preferences in mutational order in our pathway analysis (**Fig. 2d**), confirming that the accessibility of mutational trajectories – and thus, the likelihood of mutations occurring in a particular order – is indeed shaped by epistasis. Further, the differences in preferred mutational order across the affinity-only, antigen-capture, and competitive-capture models can be explained by differences in the additive and epistatic effects of mutations across different phenotypes – particularly I51Y in the HCDR2 loop, which improves affinity and polyspecificity but compromises expression (**Fig. 3b**).

**Figure 5.**
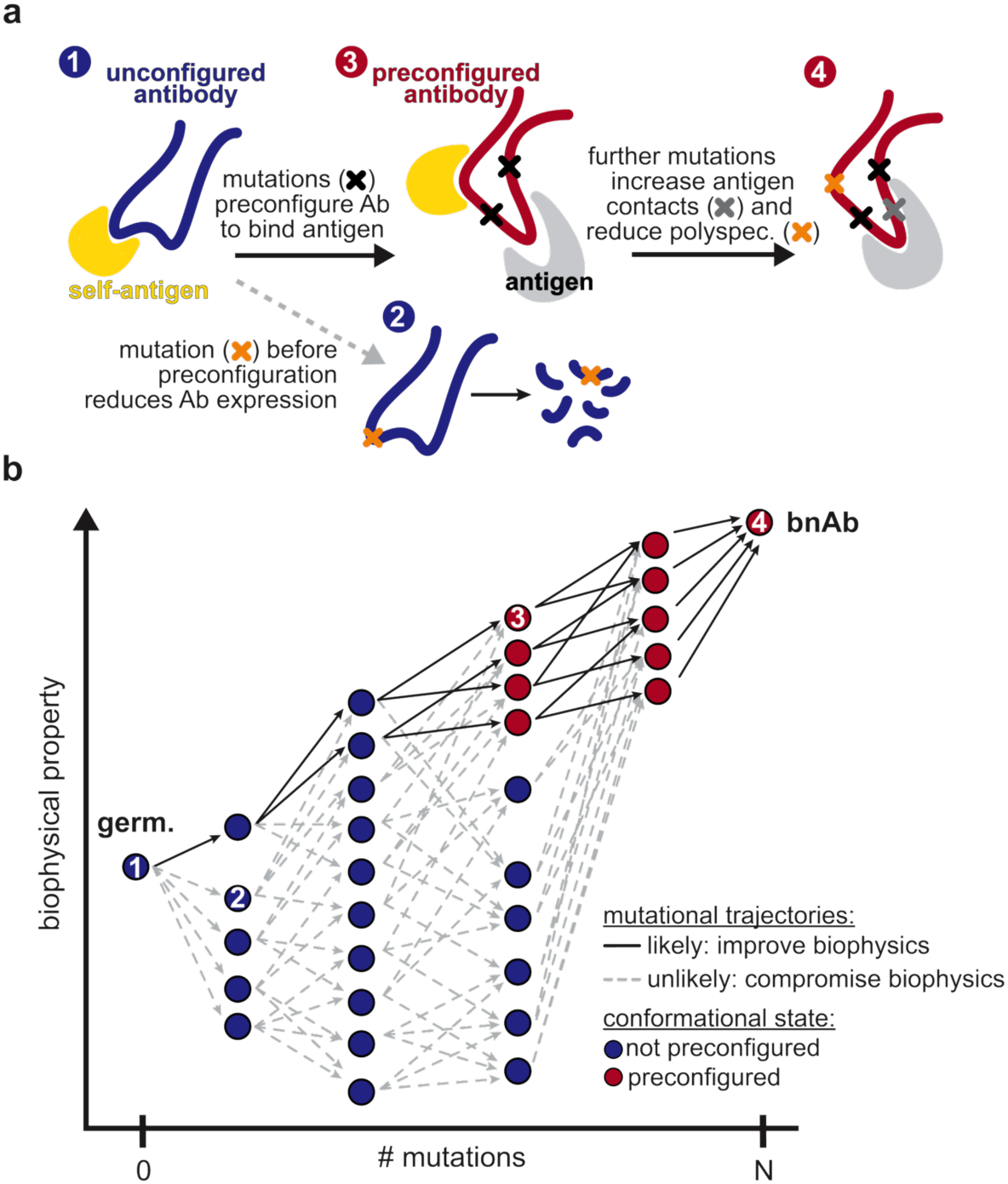
| Integrated model linking biophysical constraints, epistasis, and conformational change. **a,** Biophysical effects of mutations depend on antibody conformational state. If mutations precede antibody configuration (mediated by black “x” mutations), they often have deleterious biophysical effects (*e.g.,* on expression, orange “x”). If those same mutations follow preconfiguration, they can improve antibody biophysics (*e.g.,* polyspecificity), whereas others (grey “x”) can improve affinity through antigen contacts. **b**, Impact of antibody conformation and biophysical constraints on pathway accessibility. Most mutations compromise biophysical properties; paths containing those mutations are inaccessible (grey dotted lines). The few paths that are accessible (black solid lines) acquire mutations in a specific order that circumvents these constraints, where mutations that follow preconfiguration (red circles) have fewer biophysical costs than those that precede it (blue circles). Mutational effects 1-4 from (a) are labeled on trajectories.

To dissect epistatic interactions within the HCDR2 loop, we examined the effect of I51Y in the presence and absence of the other loop mutations, S56G and Y59F. For affinity to each antigen, we find that I51Y is most beneficial when the other loop mutations are absent and less so when one loop mutation is present (particularly with S56G), exhibiting a pattern of diminishing returns epistasis, where beneficial mutations have smaller effects in higher-affinity sequence backgrounds^58^ (**Fig. 3c, ED Fig. 7**). For expression, we observe a different pattern, where I51Y is strongly deleterious in the absence of the other loop mutations; this cost is essentially eliminated if either S56G or Y59F is present. Finally, I51Y dramatically improves (reduces) polyspecificity; this effect is consistent across sequence backgrounds but is strongest if the other loop mutations are absent. Collectively, these epistatic and pleiotropic effects explain the preferred mutational order in our pathway analysis: In the affinity-only model, I51Y is likely to occur early, as it can improve affinity on its own; in the antigen-capture model, S56G and Y59F are likely to precede I51Y to compensate for its deleterious effects on expression; in the competitive-capture model, I51Y is most likely at intermediate steps of maturation, because its deleterious effects on expression can be overcome by the benefit it has on affinity and polyspecificity.

Beyond HCDR2, we also observe strong epistasis for S31N, which neighbors the HCDR2 loop, and S94T in the LCDR3 loop. Interestingly, both mutations are highly epistatic with the HCDR2 loop mutations. S31N reduces BA1 affinity on backgrounds lacking the HCDR2 mutations but is otherwise neutral (**Fig. 3c**), reflecting its late preference in the pathway analysis (**Fig. 2d**). Conversely, S94T substantially improves affinity to BA1 on backgrounds lacking the HCDR2 mutations and is nearly neutral on backgrounds with those mutations (**Fig. 3c**), reflecting its early preference in the pathway analysis (**Fig. 2d**). Though both mutations also exhibit epistasis for expression and polyspecificity (**Fig. 3b**), those effects are outweighed by affinity in the pathway inference.

Together, these analyses suggest that mutational trajectories to Omi32 vary in biophysical accessibility – where exceedingly few trajectories are uphill – because the effects of mutations on affinity, expression, and polyspecificity are epistatic.

### Conformational rearrangement resolves steric conflicts and explains epistatic effects

To examine the biophysical basis for this epistasis, we looked to the existing Omi32-BA1 spike structure^44^. We first examined the relationship between epistatic effects and the distance between mutations, finding that, in general, the magnitude of epistasis declines with distance (**Fig. 4a**, **ED Fig. 8a**). However, there is also strong epistasis between residues that are too far apart to physically interact (> 10 Å). To assess whether this epistasis might instead be mediated by contacts with the antigen – where the effects of mutations are interdependent because they alter antigen interactions in some non-additive way – we compared the total effect of each mutation (sum of additive and epistatic effects) to the contact surface area with antigen. This analysis revealed that although some strong-effect mutations substantially contact the antigen (*e.g.,* S94T), the two strongest-effect mutations (I51Y and S56G in the heavy-chain CDR2 loop) do not (**Fig. 4b**, **ED Fig. 8b**).

To further investigate the structural basis of these strong-effect mutations and their epistatic interactions, we determined single-particle cryo-electron microscopy (cryoEM) structures of the germline and Omi32 fabs, both free and in complex with BA4 RBD (**ED Fig. 9**). To complement the existing Omi32-BA1 structure^44^, we also determined the structure for the germline in complex with BA1. We first compared interactions with each antibody to BA1 versus BA4, finding that both germline and Omi32 adopt similar conformations to bind these divergent antigens, accommodating the L452R and R493Q mutations that differentiate them (**ED Fig. 8c**)^48^. We then compared germline to Omi32, finding that despite substantial affinity differences (germline: BA1 50nM; BA4: ≥ 100 nM; Omi32: BA1 < 1 nM, BA4 ∼1 nM), these antibodies adopt similar binding conformations to each other and make comparable intermolecular contacts with both antigens (**ED Fig. 8d-e**). This unexpected similarity indicates that the affinity improvement is not driven by newly formed contacts but instead reflects differences in the conformational ensembles of the unbound antibodies.

Structures of the free fabs revealed a pronounced rearrangement of the HCDR2 and LCDR1 loops in Omi32, which are preconfigured into the antigen-bound conformation, whereas the germline loops sample an alternative state (**Fig. 4c-d, ED Fig. 10**). Though the germline can bind BA1 without rearranging the LCDR1 loops (**Fig. 4d**), the HCDR2 rearrangement occurs upon binding BA1 and BA4 (**Fig. 4c,d**). Thus, the preorganization of Omi32 likely reduces the entropic cost of binding, providing a structural mechanism for its improved affinity for antigenically divergent variants.

Interestingly, many of the mutations exhibiting the strongest epistatic effects – including I51Y, S56G, and Y59F in the HCDR2 – are in the loops that change conformation. As described above, these mutations have interdependent effects on affinity, expression, and polyspecificity (**Fig. 3c**). The structures clarify this context-dependence: I51Y would sterically clash in the germline loop configuration but is accommodated once the loop has undergone its conformational shift (**Fig. 4c**, inlay), suggesting that these mutations coordinate the loop rearrangement, and their epistasis reflects differences in the underlying loop conformational ensemble across genetic backgrounds.

The Y33F mutation in LCDR1, which also exhibits a distinct conformation in the germline compared with Omi32, as well as S94T in the neighboring LCDR3, are involved in many epistatic interactions, primarily with HCDR2 mutations (**ED Fig. 5-6**). Because each of these loops contacts the antigen, this long-range epistasis is likely mediated by conformational changes in HCDR2 and LCDR1 that affect antigen affinity.

Collectively, because many epistatic interactions involve residues that participate in this conformational transition, the accessibility of mutational pathways for this lineage is tightly coupled to loop rearrangement. These structures therefore provide a molecular mechanism for the widespread epistasis in Omi32 and reveal how conformation-mediated epistasis can shape the biophysical accessibility of antibody evolutionary routes (**Fig. 5**).

## Discussion

Here, we developed a high-throughput method for biophysically characterizing full-length natively synthesized antibodies, enabling us to examine the impact of multi-dimensional biophysical constraints on antibody affinity maturation. In applying this method to a SARS-CoV-2 bnAb, we find that evolutionary paths towards antibody breadth are highly restricted: mutations only improve affinity (and hence, breadth) if they occur in a particular order due to epistatic effects on affinity, expression, and polyspecificity. By solving the structures of the germline and mature antibodies, we find that several large-effect mutations, including highly epistatic ones, do not contact the antigen. Rather, these mutations reside in loops that undergo substantial conformational changes, thereby preconfiguring Omi32 for antigen binding. Together, these findings suggest that conformation-mediated epistasis can enable antibodies to circumvent multidimensional biophysical trade-offs.

Importantly, there are several caveats to our findings. First, we consider only the 13 mutations in the Omi32 bnAb sequence and cannot exclude the possibility that alternative, less biophysically restricted mutational paths may yield similar levels of antigen affinity and breadth. Second, BioPhy-Seq assays full-length IgGs expressed on human embryonic kidney cells, rather than B-cells. Still, we do not expect cell type to affect antigen affinity or self-reactivity, and our data suggest that expression changes are consistent across cell types. Third, our polyspecificity measurements detect nonspecific binding to a pool of human proteins that likely differs from the self-molecules present in germinal centers. Thus, these measurements capture nonspecific interactions (using a well-established reagent in the antibody field) that are typically mediated by hydrophobic effects rather than specific interactions with human proteins^8,24,59^. Finally, our models of selection neglect many of the complexities of germinal centers (*e.g.,* T-cell help, spatial structure, co-occurring mutations)^53^. Capturing these aspects in quantitative models remains an active field of research^52,60^; by contrast, our models are designed to assess the potential for antibody biophysics to shape affinity maturation.

Despite these caveats, our findings substantially advance our biophysical understanding of antibody affinity maturation. Previous work in characterizing antibody sequence-phenotype landscapes had been limited to examining antigen affinity and breadth^11,33,55,56,61–64^. Consistent with this work, many of those studies also observe strong, widespread epistasis that can shape the mutational trajectories of diverse antibodies, suggesting that epistasis is pervasive in antibody affinity maturation^55,56,62,65,66^. Notably, however, these studies observe distinct patterns of epistasis. Several studies observe synergistic epistasis, where mutations dramatically improve affinity if they occur together, but individually have undetectable effects^55,56,65,66^. In contrast, we observe that affinity-enhancing mutations have the largest effects independently, exhibiting diminishing returns when they occur together, a pattern observed in other antibodies^62,65^ and many other proteins^58,67^. Examining the molecular basis for these distinct forms of epistasis, as we do here, may enable prediction of epistasis across diverse antibody sequences.

Importantly, BioPhy-Seq enables us to measure biophysical properties beyond affinity and breadth, namely, antibody expression and polyspecificity, which are known to change in response to somatic hypermutation^8,28^ and are under selection during affinity maturation^7,9,12^. In doing so for the Omi32 bnAb lineage, we find that affinity-enhancing mutations generally reduce polyspecificity, suggesting that improvements in affinity are coupled to improvements in specificity, or alternatively, that both properties are under strong selection during affinity maturation. In contrast, we find that many sequences with improved affinity have reduced expression. While this trade-off reduces the total number of uphill paths, some paths can circumvent it by acquiring compensatory mutations that eliminate the deleterious effects of others. Our structural analysis shows that this epistasis corresponds to a conformational change in which reorientation of the HCDR2 and LCDR1 loops in Omi32 relieves potential steric clashes and preconfigures it for antigen binding, thereby improving affinity, reducing polyspecificity, and alleviating deleterious effects on expression.

To our knowledge, these are the largest quantitative biophysical datasets for full-length human antibodies. Compared with qualitative measures of affinity, expression, and polyspecificity (*e.g.,* enrichment-based assays)^24,47,61,68^, the uniquely quantitative nature of our data can be leveraged to substantially improve computational predictions of these properties^69–74^. Importantly, the epistasis observed in Omi32, along with that observed in other antibodies, cautions against using single-mutant deep mutational scanning data for training predictive models. Because it is infeasible to generate combinatorial datasets for all human antibody sequences, we instead advocate for examining the combined effects of mutations across divergent antibody sequences to uncover the general nature and magnitude of epistasis. By improving the accuracy of predictive models, this work will advance efforts to engineer antibodies that bind targets of interest and exhibit desirable clinical characteristics (*e.g.,* low polyspecificity and high expression). Further, such predictive tools will enable mapping biophysical properties across human antibody repertoires, advancing our fundamental understanding of affinity maturation and informing the design of vaccine antigens that elicit antibodies with favorable biophysical parameters (*e.g.,* mutational paths to bind some epitopes may be less biophysically constrained than others). Beyond improving predictions for antibodies, the adaptability of BioPhy-Seq will enable similarly large, high-resolution biophysical datasets and, consequently, accurate computational predictions of biophysical properties for other human membrane proteins.

Further, compared with other proteins, structural predictions for antibodies remain challenging and are of major interest for predicting binding interactions^75–78^. Our structural data show that, due to mutation-induced changes in antibody conformation, antibody-antigen structures are not necessarily predictive of binding affinity^69,79^. Because exceedingly few germline antibody structures exist in the Protein Data Bank (a tiny fraction of which are unbound)^80^, the extent to which affinity maturation causes preconfiguration remains unknown^66,81–84^. Given the implications of preconfiguration for shaping the breadth and specificity of human antibody repertoires, this work motivates further structural work to discern the effects of somatic hypermutation and antigen binding on the conformations of diverse antibody sequences.

Together, this work leverages BioPhy-Seq, a new experimental platform for biophysically characterizing natively synthesized full-length antibodies, to advance our fundamental understanding of how mutations alter antibody biophysics. In future studies, we envision applying BioPhy-Seq to divergent antibody sequences that mature to recognize distinct molecules, enabling us to define the general biophysical mechanisms that shape antibody maturation and to harness them to engineer antibody therapeutics and predict immune responses. Further, the adaptability of BioPhy-Seq enables its application to diverse human membrane proteins, thereby facilitating the discovery of generalizable biophysical mechanisms underlying protein variation and evolution.

## Acknowledgements

We would like to acknowledge Kyle Lucier, Skerdi Senko, Nhi Tran, Ashlyn Farwell for cryoEM grid preparation and imaging, Jimin Lee for initiating experiments for this project, Kenneth Matreyek and Nicholas Wu for sharing HEK landing pad cells, Timothy Tan for sharing culturing and integration protocols, Christopher Barnes for sharing the recombinant spike expression plasmids, Peter Rowheder and Charles Craik for assistance with making BLI measurements, and Daniel Maurer, Christopher Barnes, Matthew Shoulders, James Chen, and members of the Phillips lab for their thoughtful comments,. Research reported in this publication was supported by the Howard Hughes Medical Institute (to AP), the National Institute of Allergy and Infectious Diseases of the National Institutes of Health under Award Number R01AI189532 (to AP), the UCSF Discovery Fellows Program (to CT and SG), and the National Science Foundation (GRFP awarded to AK). Sequencing was performed at the UCSF CAT, supported by UCSF PBBR, RRP IMIA, and NIH 1S10OD028511-01 grants. Flow cytometry was performed at the PFCC (RRID:*SCR_018206*), supported in part by Grant NIH P30 DK063720 and by the NIH S10 Instrumentation Grant S10 1S10OD021822-01, as well as technical support from Vinh Nguyen and Priscilla Sanchez.

## Methods

### Antibody sequences and mutations of interest

The germline sequence (**SI File 1**) was inferred from the Omi32 sequence (**SI File 2**) using IgBlast^44,85^. These sequences differed by thirteen amino acid substitutions, all of which were included in the library, using the codons in the Omi32 nucleotide sequence (*i.e.,* no codon optimization was performed).

### IgG antibody construct and cell line generation

The IgG construct is annotated in **SI Files 1-2** and is comprised of: a signal peptide, the variable and constant domains of the heavy chain with a synthetic transmembrane domain (from platelet derived growth factor), a P2A cleavage site, a second signal peptide, the light chain variable and constant domains, and a C-terminal myc-epitope tag – resulting in a membrane-bound IgG with an epitope tag on the light chain. This construct was cloned into a vector containing *attb* recombination sites (for integration into a genomic landing pad) and a downstream internal ribosomal entry site (IRES) followed by a puromycin resistance cassette.

To generate IgG-expressing cells, the entire *attb* plasmid was integrated into a Tet-inducible genomic landing pad (BFP-2A-iCasp9-2A-Blast^R^) in Human Embryonic Kidney (HEK) cells that co-express rtTA and the Bxb1 recombinase, as previously described^42^. Briefly, landing pad cells were maintained in high-glucose DMEM (Thermo 11995073) with 10% FBS (Corning 35-010-CV), 1% penicillin-streptomycin (Thermo 15140122), 1% non-essential amino acids (Fisher 11-140-050), 1% GlutaMAX (Fisher 35-050-061), and 2 μg/mL doxycycline (Fisher AAJ67043AD). To integrate the IgG construct, 6 x 10^5^ cells were seeded in 6-well plates and transfected with 1.2 μg attb plasmid and 5 μL Fugene 6 (Promega E2693), following the manufacturer’s instructions (day 1). On day 2, 500 μL of complete growth media was added dropwise to the cells. Negative selection was performed on day 4 by replacing the media with negative-selection media (complete growth media supplemented with 2 μg/mL doxycycline and 10 nM AP1903 (MedChem Express HY-16046)). On day 5, cells were allowed to recover by discarding the negative selection media, washing once with sterile 1X PBS (Fisher MT21040CV), and replacing with 2 mL complete growth media supplemented with doxycycline. On day 7, cells were declumped by washing with PBS, trypsinizing, and resuspending in 2 mL complete growth media containing doxycycline. Positive selection was initiated on day 8 by replacing the growth media with fresh growth media containing doxycycline and 1 μg/mL puromycin (Thermo A1113803). Cells were maintained in positive selection media for at least 7 days before performing flow cytometry, subculturing as needed.

### Combinatorial Golden Gate Assembly and Library Cloning

The 13 mutations were partitioned across 8 fragments (3 fragments for the heavy chain variable region, 3 fragments for the light chain variable regions, 1 fragment for the heavy chain constant regions, and 1 fragment for the light chain constant regions) with each variable chain fragment containing 1-3 mutations such that the combinatorially complete 2^13^ library could be constructed from 34 fragments (fragment 1: 2^2^, fragment 2: 2^3^, fragment 3: 2^1^, fragment 4: heavy chain constant region, fragment 5: 2^3^, fragment 6: 2^3^, fragment 7: 2^1^, and fragment 8: light chain constant region). Fragments were designed as described previously^55^, each flanked by BsmbI (New England Biolabs #R0739L) sites and unique 4-bp overlaps to ensure correct assembly and were generated by PCR using Q5 polymerase (New England Biolabs #M0491L; primers for generating fragments are listed in **SI File 3**). Amplified fragments were purified using beads (Aline Biosciences #C-1003-50), and concentrations were measured by AccuGreen (BioTium 31066). Fragments were pooled for assembly by combining all versions of the same fragment in equimolar ratios and subsequently combining fragment pools at equimolar ratios, generating a fragment mixture that, once assembled with a 16N barcode (added to the light chain constant region with a primer; **SI File 3**) and the *attb* plasmid in a single Golden Gate reaction, produced all 2^13^ antibody sequences in approximately equal frequencies^86^. Assemblies were purified by column clean-up (GeneAid PDH300), eluted with water, and transformed into electrocompetent DH10B (NEB 3020), per the manufacturer’s instructions. Following recovery, transformed cells were suspended in semi-solid 0.3% agarose (SeaPrep Agarose; Lonza 50302) in LB (1% tryptone, 0.5% yeast extract, 1% NaCl): before adding cells, solution was heated to dissolve agarose; once agarose cooled to ∼30°C, 50-500 μL transformed cells were suspended into separate 200 mL agarose solutions in 1 L baffled flasks and ampicillin (VWR 76344-932) was added to a final concentration of 100 μg/mL; flasks were moved to cold room for 2 hours to allow the agarose to solidify; flasks were then gently moved to 37°C (no shaking) to allow bacterial colonies to grow in the agarose suspension. Dilutions of the transformation were also plated on LB-Amp plates to estimate transformation efficiency and the number of cells per flask. After 16-20 hours of growth, LB-amp plates were counted to select the flask with the optimal number of colony-forming units (CFU, see below), and that flask was shaken at 37°C for 1 min to homogenize the bacterial colonies in the agarose. To increase DNA yield for subsequent plasmid library transfection while maintaining even library diversity, 40 mL of the homogenized bacterial colonies were transferred to a new 1L baffled flask and diluted to a final volume of 200 mL with LB and final concentration of 100 μg/mL ampicillin. Cells were shaken for approximately 5 hours, with optical density (OD) readings taken every 30 minutes, such that each cell replicated approximately 3-4 times. Cells were pelleted by centrifugation (3,000 x g, 10 min) and plasmid library was midiprepped for downstream transfection (Zymo Research #D4201). A flask containing approximately 250,000 CFU was chosen to generate the barcoded antibody library, exceeding the antibody sequence diversity (8,192) but constraining it so that all plasmids would be sequenced (at ≥3X coverage) on a single PacBio flow cell.

### Long-read sequencing to map barcodes to antibody sequences

Plasmid library was prepared for PacBio sequencing by restriction digestion with BsaI (New England Biolabs #R3733L) to produce a 3,600 bp DNA fragment spanning the full IgG construct and 16N barcode sequence. The 3,600 bp fragment was isolated by 1% agarose gel electrophoresis with SYBR Safe DNA Gel Stain (Invitrogen #S33102) on a blue light transilluminator to mitigate DNA damage. The band of interest was purified with a Monarch Spin DNA Gel Extraction Kit (New England Biolabs #T1120S) and eluted in Qiagen Buffer, EB (Qiagen #19086). The fragment library was sent to the UC Davis DNA Technologies and Expression Analysis Core Facility for PacBio sequencing library preparation and long-read sequencing on a PacBio Revio at >30x coverage per barcoded antibody sequence.

Barcodes were associated with antibody sequences using the dms-vep-pipeline-3 Github repository^87^ (https://github.com/dms-vep/dms-vep-pipeline-3). Briefly, PacBio circular consensus sequences (CCSs) were aligned to the IgG construct using *alignparse*, and consensus sequences for each antibody-barcode pair were determined, requiring at least two consensus sequences per barcode. Custom code was then used to assign binary genotypes to each antibody variant-barcode pair and export the variant-barcode table for downstream Illumina sequencing analysis, where any IgG constructs with non-designed (*i.e.,* spurious) mutations or indels were removed from the dataset, resulting in ∼125,000 high-confidence and unique IgG-barcode pairs (see Data and code availability below).

### HEK antibody library production

Plasmid library was transfected and integrated into HEK landing pad cells as described above, scaling to 50 million cells to maintain ∼125,000 uniquely barcoded IgG, minimize bottlenecking of low-diversity variants, and generate the cell library in a single transfection. Following negative and positive selection (as described above), the HEK antibody library was expanded for two passages, and cells were frozen in aliquots of 10 million cells at each passage. Aliquots were subsequently thawed and used immediately for each replicate BioPhy-Seq experiment after one post-thaw recovery passage.

### SARS-CoV-2 spike receptor binding domain (RBD) antigen

Biotinylated SARS-CoV-2 spike RBD was purchased from AcroBiosystems (Wuhan: SPD-C82E9; BA1: SPD-C82E4; BA4/5: SPD-C82EW). Wuhan spike RBD was reconstituted in 42 μL (21 μM), BA1 spike RBD was reconstituted in 125 μL (7 μM), and BA4 spike RBD was reconstituted in 125 μL (7 μM) according to the manufacturer’s recommendation. All solutions were aliquoted, snap-frozen in liquid nitrogen, and stored at -80°C until use.

### Polyspecificity reagent (PSR)

Polyspecificity reagent was prepared as described previously^24^. Briefly, two billion HEK Freestyle cells (ThermoFisher #R79007) were pelleted (550 x g, 3 min) and resuspended with 200 mL Buffer A (50 mM HEPES, 150 mM NaCl, 2 mM CaCl_2_, 5 mM KCl, and 5 mM MgCl_2_, pH 7.2). After a second spin, cells were resuspended in 60 mL of Buffer B (50 mM HEPES, 150 mM NaCl, 2 mM CaCl_2_, 5 mM KCl, 5 mM MgCl_2_, and 10% glycerol, pH 7.2). Cells were pelleted a third time and resuspended in three pellet volumes (approximately 6 mL) of Buffer B with freshly dissolved Pierce Protease Inhibitor Mini Tablets (1 tablet per 4 mL of Buffer B, Thermo Scientific #A32961). After resuspension, cells were sonicated with a probe sonicator set at 8W output for 30-second pulses on ice until no live cells could be viewed under a microscope (brightfield), resulting in ∼8 total pulses. Lysed cells were then pelleted (2,100 x g for 5 min), supernatant was collected, and total protein concentration was assessed using a Pierce 660nm Protein Assay Reagent (Thermo Scientific #1861426) and Albumin Standards (Thermo Scientific #23209). The N-terminal amines of soluble cytosolic proteins (SCPs) in the supernatant were biotinylated using 1 mg of EZ-Link Sulfo-NHS-Biotin (Thermo Scientific #21217) per 9 mg of protein by adding the requisite volume of 10 mM Sulfo-NHS-Biotin to the supernatant and rotating the mixture for 16-20 hours at 4°C. The solution was passed through PD-10 gravity desalting columns (Cytiva #17085101), equilibrated with Buffer A, to remove excess Sulfo-NHS-Biotin and to exchange buffers. The biotinylated soluble cytosolic protein (b-SCP) solution was then quantified using the Pierce 660nm Protein Assay Reagent and prediluted Albumin Standards to a final protein concentration of ∼8.5 mg/mL, flash-frozen in 1 mL single-use aliquots, and stored at -80°C until use.

### BioPhy-Seq assays

Each BioPhy-Seq measurement was performed in biological duplicate (except expression, which was measured in 6 biological replicates), on different days. Except where noted, cells were pelleted by spinning at 300 x g for 5 minutes.

#### Cell labeling – affinity and expression measurements

For each sample, 20 million cells displaying the Omi32 IgG library were harvested using versene (ThermoFisher #15040066), washed twice with cold FACS buffer (clear DMEM (ThermoFisher #21063029), 2% FBS (Corning 35-010-CV), 0.5 mM EDTA (Invitrogen #AM9260G)), counted, resuspended in RBD diluted to the desired concentration in FACS buffer (spanning 100 nM to 1 pM, in 1-log increments), and incubated for 20 hours (to reach equilibrium) at 4°C with gentle rocking. The total volume of the labeling reaction was adjusted to limit ligand depletion effects to <10%, ranging from 50 μL for 100 nM RBD to 1 L for 10 pM RBD. Following labeling with RBD, cells were pelleted by centrifugation at 4°C (for large volumes, cells were spun at 300g for 10 min; smaller volumes were spun for 5 min), washed twice with FACS buffer and stained with streptavidin-PE (1:50; ThermoFisher S866) and myc-FITC (1:50; Miltenyi Biotec 130-116-485) by incubating for 1 hour at 4°C in the dark with gentle rocking. Cells were then washed twice with FACS buffer and resuspended in FACS buffer to a concentration of 5-10 million cells/mL for downstream flow cytometry.

#### Cell labeling – polyspecificity measurements

Cells were prepared as described above but were labeled with polyspecificity reagent (PSR) instead of RBD and were fixed following secondary labeling and two washes with FACS buffer by treating with 4% PFA (Biotium #22023) for 10 minutes. Fixing was performed for these measurements to minimize dissociation during sorting. After fixing, cells were washed twice with FACS buffer and resuspended in FACS buffer to a concentration of 5-10 million cells/mL for downstream flow cytometry.

#### Sorting and recovery

For each sample, 1-2 million cells were sorted into four tubes containing 200 uL ice-cold PBS to ensure adequate library coverage (100x). For expression, single cells were sorted into four, six, or eight populations based on FITC fluorescence intensity, with each gate capturing 25%, 16.7%, or 12.5% of the library, respectively. For binding and polyspecificity, single IgG+ cells were sorted based on PE fluorescence using four gates – one containing non-binders and the other three containing 33% of the remaining population (low, medium, and high PE intensity). Sorting scheme is sketched in **SI File 5**. Sorted cells were pelleted (300 x g for 1 min) and left in FACS tubes overnight at 4°C for DNA extraction the following day.

#### DNA extraction, library preparation, and sequencing

For unfixed sorted cells (affinity and expression measurements), genomic DNA was extracted using the Qiagen DNeasy Blood & Tissue Kit (Qiagen #69504), following the manufacturer’s instructions with the addition of a 5-minute room-temperature incubation with 4 μL 100 mg/mL RNase A (Qiagen #19101) before adding Buffer AL, and eluting in 50 μL of double-deionized water. Eluent was added back to the column for a second elution to increase DNA yield. For fixed sorted cells, genomic DNA was extracted using phenol-chloroform extraction. Briefly, fixed cells were pelleted (500 x g for 3 min) in FACS tubes, resuspended in 400 μL lysis buffer (1% SDS, 50 mM Tris-HCl, 10 mM EDTA, pH 8) with 16 μL of 5M NaCl per 5 million cells (scaling accordingly for samples with more cells). Each sample was incubated for 16-20 hours at 66°C. After incubation, 8 μL of 20 mg/mL Proteinase K (Qiagen #RP101B) was added to the mixture, cells were briefly vortexed and then incubated at 55°C for 1 hour. Samples were loaded into phase-lock tubes with 400 μL Phenol:Chloroform:Isoamyl Alcohol (25:24:1). Phase-lock tubes were made by spinning 200 μL DuPont MOLYKOTE High-Vacuum grease (VWR #100504-358) at 10,000 x g for 1 min in 1.5 mL microfuge tubes. After addition to phase-lock tubes, samples were vortexed and spun (20,000 x g for 5 min). The aqueous (top) phase was transferred to low-bind 1.5 mL microfuge tubes (Sorenson Bioscience #39640T) and DNA was precipitated with isopropanol, 40 μL of 3M Sodium Acetate, and 1 μL GlycoBlue (Invitrogen #AM9515) and incubated at -80°C for 1 hour. DNA was pelleted (20,000 x g for 5 min), washed with 1 mL of 80% ethanol, and pelleted again. After removing ethanol and allowing the pellet to air dry, it was resuspended in double-deionized water and incubated at 65° for 15 minutes.

Sequencing libraries were prepared by performing a two-stage PCR as previously described^55^– the first stage appended unique molecular identifiers (UMI, to account for biased PCR amplification), sample indices (for multiplexing and to ensure sequence diversity on the flow cell), and a partial Illumina adapter; the second stage appended the rest of the Illumina adapter and N7 and S5 Illumina sample indices (for additional multiplexing; sequencing library preparation primers in **SI File 4**). In the first stage, 500 ng of gDNA was amplified with 50 μL PCR reactions using PrimeStar GXL DNA polymerase (Takara Bio, #R050A) following the manufacturer’s instructions (95°C for 3 min, 98°C for 10 s, 68°C for 3 min and 15 s, repeat 2-4 14x, 68°C for 3 min) with each primer at 0.25 μM; 2-20 reactions were performed for each sample such that all gDNA was used to ensure adequate library sampling. PCR product volume was increased to 100 μL with double-deionized water and was purified using 60 μL beads (Aline Biosciences #C-1003-50), eluting in 50 μL double-deionized water. Purified PCR products from the same sample were pooled and mixed well. For the second stage PCR, 20 μL of purified PCR product was amplified in 50 μL PCR reactions using PrimeStar GXL DNA polymerase according to the manufacturer’s instructions (95°C for 3 min, 98°C for 10 s, 66°C for 15 s, 68°C for 1 min, repeat 2-4 29x, 68°C for 3 min) with each primer at 0.25 μM; multiple PCRs were performed per sample such that 20% of the total first stage PCR product was used (again to maintain library diversity). PCR product volume was increased to 100 μL and purified again using a double-sided bead purification, with 70 μL of beads added to remove high molecular weight contaminants and 30 μL of beads were then added to isolate the final product from low molecular weight contaminants with a final elution in 50 μL of double-deionized water. Purified PCR products from the same sample were pooled and mixed well. Sequencing library concentration was determined by Accugreen (BioTium 31066). For each sorted population, samples were pooled based on cell count, and subsequently pooled in equimolar ratios. The size of the fragment pool was verified to be 328 bp by HS DNA1000 TapeStation (Agilent 5067-5584). Samples were sequenced on an Illumina NovaSeqX using 150bp paired-end reads, ensuring >10 million reads per sample (>400-fold coverage per variant).

#### Sequencing data processing

Reads were demultiplexed first by N7 and S5 indices. Inline indices, UMIs, and variant barcodes were extracted from the resulting sequences using a custom parsing script (see Data and code availability below). Reads with mismatched inline indices were discarded, UMIs were deduplicated per barcode to account for PCR bias, deduplicated barcodes were converted to binary genotypes, and binary genotypes were counted for downstream analyses. Sequenced barcodes were required to be an exact match with the variant-barcode table and were excluded from analysis if no match was found.

### Tite-Seq *KD* inference

Binding affinities were inferred using the mean-bin approach, as described previously^33,47,55^. Briefly, sequencing data (the number of counts of each variant sequence *s* in bin *b* at concentration *c*, *R_b,s,c_*) were integrated with flow cytometry data (the mean, *F*, and standard deviation, *σF*, of the fluorescence intensity for sorted cells in each bin *b* at each concentration *c*, and the number of cells, *C*, sorted into each bin *b* at each concentration *c*). First, the mean log-fluorescence for each variant sequence at each concentration was calculated as:

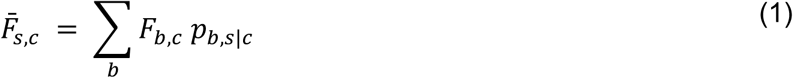

Where *p_b,s|c_* is the probability a cell with sequence *s* is sorted into bin *b* at concentration *c*, and is estimated from the sequencing data as the fraction of reads in that bin corresponding to sequence *s*:

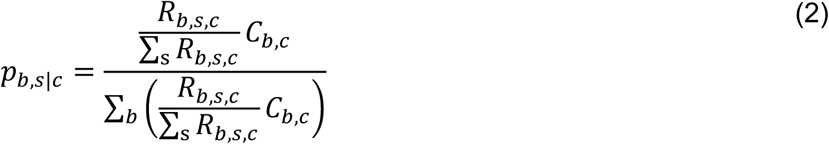

The uncertainty is propagated as:

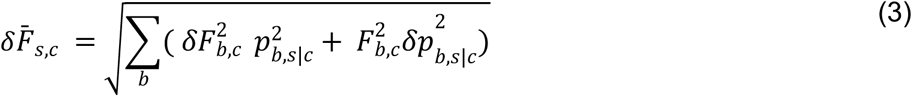

Where 𝛿*F_b,c_* is the spread in log-fluorescence of a specific bin. These distributions are not uniform, but we find that the standard deviation, *σF_b,c_*, captures the variation across bins. The error in *p_b,s|c_* largely arises from sampling biases during sequencing, which we approximate as a Poisson process, giving:

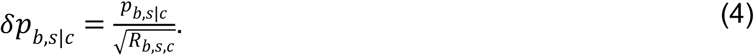

Equilibrium binding affinities (*K_D_*) are then inferred for each variant by fitting the Hill function across the seven antigen concentrations *c* (M):

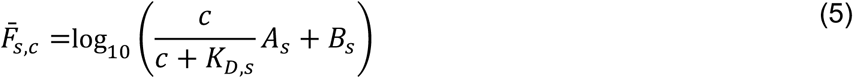

where *A_s_* is the increase in fluorescence due to saturation, and *B_s_* is the background fluorescence. Fitting was performed using the *scipy.optimize curve_fit* function. Inferred *K_D_* outside the titration boundaries were pinned to the boundaries (10^-7^ and 10^-11^ M). Inferred *K_D_* were quality-controlled using the following criteria (detailed in the GitHub repository below). First, antibody variants with low log_10_(*A_s_*/expression) relative to the median of each dataset were classified as false-binders and pinned to the upper detection boundary (10^-7^ M). These pinned values were retained in the dataset if the variant had sufficient sequencing support (>10^-7^ frequency) and its biological replicate was within 1 -log_10_*K_D_* unit of the lower detection boundary (≤10^-8^ M). If those criteria were not met, that variant was removed from the dataset.

Second, antibody variants with high log_10_(*A_s_*/expression) relative to the median of each dataset were required to satisfy goodness-of-fit criteria (R^2^ ≥ 0.8 and residual standard error ≤ 1.0). Only variants with both biological replicates passing all of these criteria were retained for downstream analysis. Mean and SEM -log_10_*K_D_* for retained variants were calculated and used for downstream analysis.

### Expression inference

Expression values were inferred as described previously^55^. Briefly, the IgG library was sorted into 4-8 bins along the FITC fluorescence intensity axis. The mean log_10_ FITC fluorescence was computed for each variant using the variant sequencing counts and fluorescence data, as described above for the *K_D_* inference (Equation 1).

To correct for run-to-run differences in fluorescence intensity (*e.g.*, variation in PMT voltage or detector gain), expression values within each replicate dataset were shifted by a constant equal to the difference between the dataset median and the median of the six replicates. For each variant, the six median-normalized log_10_ FITC fluorescence values were used to compute the mean expression and SEM. Variants with SEM ≥ 0.5 were removed after determining that these high-error measurements were associated with low sequencing coverage.

Because each flow cytometry experiment was performed under identical conditions (including saturating secondary labeling, post-thaw passaging times, detector voltages set such that fluorescence measurements remained within the linear range of the PMT), differences in log_10_ FITC fluorescence are expected to be proportional to log_10_ differences in the number of IgG copies on the surface of HEK cells. Accordingly, fluorescence differences are reported as Δlog_10_ copy #*_app_*.

### Polyspecificity inference

Polyspecificity scores (*EC_50_*) represent the concentration of polyspecificity reagent corresponding to 50% occupancy. These scores were inferred in the same way as the binding affinities described above, using the Hill equation to fit the mean fluorescence intensity (inferred from sequencing and flow cytometry data, as described above) to the concentration of the polyspecificity reagent (in mg/mL). Biological duplicates that passed goodness-of-fit criteria (R^2^ ≥ 0.8 and residual standard error ≤ 1.0) were averaged for downstream analysis.

### Isogenic validation

Low-throughput isogenic measurements were made in biological duplicate for a subset of variants to validate high-throughput BioPhy-Seq measurements. HEK cells expressing a clonal IgG sequence were generated and labeled as described above, except cell number was scaled down to 1 million cells per sample. Rather than sorting and sequencing, measurements of affinity, expression, and polyspecificity were made by analytical flow cytometry, collecting cell fluorescence intensities for PE (affinity and polyreactivity) and FITC (expression). Like the BioPhy-Seq inference, binding affinity (*K_D_*) and polyspecificity (*EC_50_*) were inferred for each variant by fitting the following function to the mean log PE fluorescence:

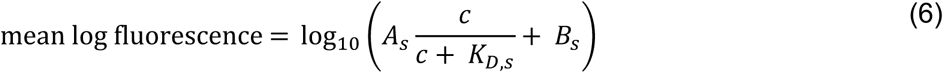

where *c* is the concentration of antigen (M) or polyspecificity reagent (mg/mL), *A_s_* is the increase in fluorescence due to saturation, and *B_s_* is the background fluorescence. Binding affinities and polyspecificities were averaged across biological replicates. For expression, rather than averaging mean log_10_ FITC fluorescence across experiments for each variant, the change in expression of each variant relative to germline was calculated for each replicate (Δlog_10_ copy #*_app_*). Replicates were averaged and compared to the corresponding Δlog_10_ copy #*_app_* inferred from the high-throughput dataset.

### Force-directed layouts

To reduce the dimensionality of the antibody sequence-biophysics landscape, we implemented force-directed layouts, as described previously^55^. Briefly, each antibody sequence is represented by a node that is connected by edges to all single-mutation neighbors. An edge between nodes *s* and *t* are weighted by the biophysical change resulting from the corresponding mutation. For the layouts presented in **Figure 2e**, BA1 affinity was used for the edge weights:

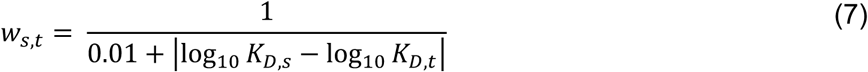

Thus, if a mutation does not impact BA1 affinity, those nodes will be close together, and vice versa. An interactive form of this graph is available as an online data browser here: https://amphilli.github.io/Omi32_browser_git/.

### Epistasis inference

#### Nonlinear transformation

As described previously^55,88^, prior to inferring mutational effects and epistatic coefficients, each biophysical property was transformed onto an additive scale. Because -log*K_D_* are proportional to free energies, which are expected to combine additively^89^, we applied linear models directly to the -log*K_D_*. For expression and polyspecificity, which do not correspond to free energies, we also proceeded with log-transformed values. Importantly, because measurements can deviate from additivity for several reasons (*e.g.,* the property or measurement is inherently non-linear, measurement noise or detection limits, *etc.*), we further transformed each log-transformed property onto an “additive” scale (as described previously^90^) and applied linear models to both the transformed and untransformed data. Models in the main text are those with the fewest parameters, highest performance, and biological interpretability; all other models are presented in **ED Fig. 4**.

#### Linear interaction models

To infer the effects of mutations, we implement linear models where the biophysical property is the sum of the effects of the individual mutations and combinations thereof. The additive model is given by:

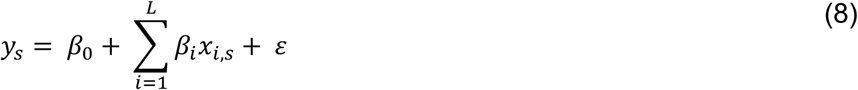

Where *L* corresponds to the 13 mutations in Omi32, *β_0_* is an intercept, *β_i_* is the effect of mutation at site *i*, *x_i,s_* is the genotype of variant sequence *s* at site *i*, and 𝜀 represents independently and identically distributed errors. The general epistatic model is thus given by:

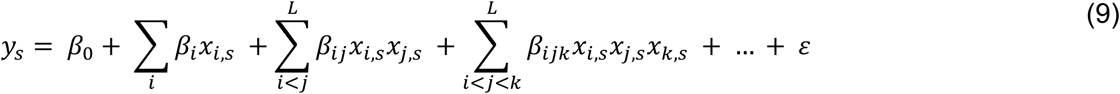

Where *β_ij_* are second-order interaction coefficients between sites *i* and *j*, *β_ijk_* are third-order interaction coefficients between sites *i*, *j*, and *k*, and so on, up to a specified maximum order of interaction.

To encode the binary genotypes *x_i,s_*, we implement two frameworks that produce identical models but yield different coefficients and thus have distinct uses. First, we use *x_i,s_* ∈ [0,1], known as a “reference-based” framework, which encodes all coefficients relative to the germline sequence. Second, we use *x_i,s_* ∈ [-1,1], known as a “reference-free” framework, which encodes all coefficients relative to the dataset average. These frameworks are described in detail elsewhere^57,88^; the reference-based framework is presented in the main text, as it is most useful for examining phenotypic effects relative to a sequence of interest, and the reference-free framework is presented in **ED Fig. 6**, as it is useful for those interested in protein sequence architecture.

Because Omi32 has 13 mutations, there are 13 possible orders of interactions, with 2^13^ possible coefficients *β*. Rather than infer a 13^th^-order model, which would have an equivalent number of measurements and parameters, we take a conservative approach that is less prone to inferring spurious higher-order terms (as described previously^88^) Specifically, we infer models truncated at each order and evaluate model performance by cross-validation, using 90% of the data to train the model and the remaining 10% to evaluate predictive performance (*R^2^*). We then average performance across 8 folds for each truncated model, determine the order that maximizes performance, and then re-train the model truncated at this optimal order using the full dataset to obtain the final coefficients. For biophysical properties where higher-order models modestly improved performance (*i.e.,* marginally above the SEM from cross-validation), model predictions were compared to low-throughput validation data to choose the optimal model and avoid over-fitting the data. For each biophysical property, we find that the number of measurements exceeds the number of coefficients by at least an order of magnitude; thus we do not implement regularization for downstream analysis.

To train these models, we perform ordinary least squares regression using the Python package *statsmodels*, which yields coefficients *β* with standard errors and *p*-values. We evaluate the statistical significance of coefficients using a *p*-value of 0.05 with a Bonferroni correction by the total number of model parameters.

#### Structural analysis of epistasis

To contextualize these epistatic effects, we performed two analyses using the previously characterized BA1-Omi32 structure (PDB ID: 7ZFE^44^). First, we used ChimeraX *buriedarea* function^91^ to compute the buried surface area between RBD and each mutated residue in Omi32, using the default radius to approximate a water molecule (1.4 Angstroms). Second, we used PyMol^92^ to measure the distance between ɑ-carbons for all possible pairs of mutated sites.

### Phenotype predictions

Epistasis models were used to predict all five phenotypes for each of the 8,192 genotypes in the library. Because many genotypes were missing data (due to measurement noise, low sequencing coverage, etc.), we used the predicted phenotypes for all figures except **Figure 3** and **ED Fig. 4-7**. These predicted phenotypes are compared to experimental phenotypes and the low-throughput validated phenotypes in **ED Fig. 1**. This comparison revealed that the predicted phenotypes are better correlated to the low-throughput measurements than the experimental values, likely because the epistatic models can remove experimental noise.

### Pathway analysis

We infer the likelihood of all possible mutational trajectories from the germline to Omi32 by extending previously described affinity models^55^ to incorporate expression and polyspecificity measurements. Importantly, irrespective of the biophysical parameters used, these models are an intentional oversimplification of selection in germinal centers – designed to capture the potential for biophysical effects to shape mutational trajectories. Incorporating the impact of non-biophysical factors (*e.g.,* spatial structure, T-cell mediated selection, *etc.*) is an active area of research and beyond the scope of this work^52,60^.

#### Selection models

We construct simple models restricted to the weak-mutation regime, in which mutations occur and are selected independently^50^; further, we do not allow mutations to revert. To estimate the fixation probability of each mutation, we extend previously developed models that were solely based on binding affinity^55^. Here we expand on these prior models to integrate BioPhy-Seq expression and polyspecificity measurements by estimating ‘antigen-capture probabilities’, which represent the combined impact of a mutation on antibody expression, polyspecificity, and antigen binding affinity, and hence the likelihood of impacting B-cell antigen-capture. These probabilities are defined from the classical fixation probability:

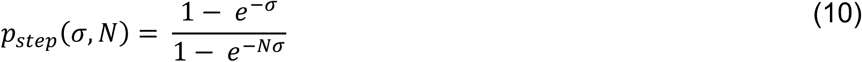

Where *p_step_* is the probability of a single mutational step for a mutation with selection coefficient *σ* in a population of size *N*^51^. Here, *σ* is proportional to the log_10_ difference in antigen-capture between sequences *s* and *t,* which we define with three separate models: an ‘affinity-only’ model (Equation 11) identical to our prior work^55^, where mutation probabilities increase with their improvement in affinity, an ‘antigen-capture’ model (Equation 12), where mutations probabilities increase with their improvement in the total number of antigen-bound antibodies on the surface of a cell, and a ‘competitive-capture’ model (Equation 13), where mutation probabilities increase with their improvement in the total number of antigen-bound antibodies on the surface of the cell, accounting for competitive interactions with host proteins. Prior to implementing these models, polyspecificity (log*EC_50_*) and expression (log*copy #_app_*) were normalized to the same scale as BA1 affinity (-log*K_D_*), enabling direct comparison of mutational effects across these properties.

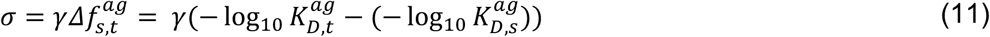

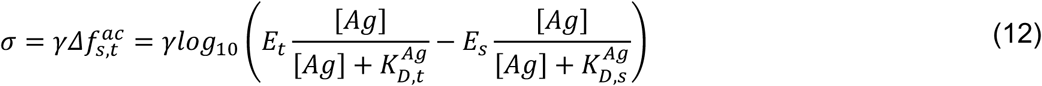

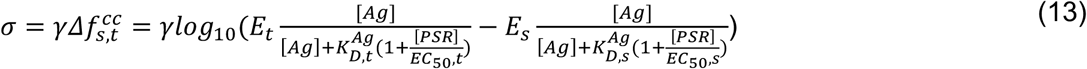

Where *E_t_* and *E_s_* are expression values of variants *t* and *s*; similarly, *EC_50,t_* and *EC_50s_* are polyspecificity values of variants *t* and *s*. Notably, the antigen-capture and competitive-capture models require additional parameters, including antigen concentration ([*Ag*]) and polyspecificity reagent concentration ([*PSR*]). Because the effective concentrations experienced by B cells in germinal centers remain poorly defined^9^, these parameters were not chosen to represent precise physiological values. Instead, we selected parameters such that the relative contributions of affinity, expression, and polyspecificity could be evaluated and compared across mutational trajectories (detailed below).

Analyses were performed in the low-antigen regime ([*Ag*] << *K_D_*), where antigen is limiting, and the number of antigen-bound receptors depends approximately on the ratio of expression to affinity. In this regime, improvements in affinity and expression contribute independently and additively (in log space) to antigen-capture:

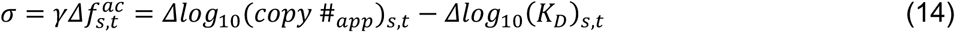

For the competitive-capture model, we considered a high-polyspecificity regime ([*PSR*] >> *EC_50_*), in which nonspecific engagement strongly competes with antigen binding. In this limit, increases in polyspecificity act as an another multiplicative penalty on antigen-capture. As a result, the effects of affinity, expression, and polyspecificity become approximately additive, yielding a simple linear approximation that makes the contribution of each biophysical parameter to selection interpretable:

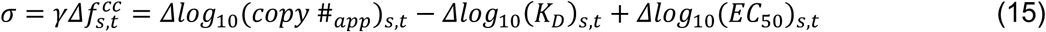

Graphical representations of these independent, log-linear relationships are shown in **ED Fig. 3**.

The strength of selection in these models can be tuned using *N* (effective population size) or 𝛾. As described previously, here we use two sets of parameters to span varying strengths of selection: (1) weak selection (*N* = 20; 𝛾 = 0.5) and (3) strong selection (*N* → ∞; 𝛾 → ∞, such that *p_step_* = 1 if Δ𝑓*_s,t_*> 0 and 0 otherwise). Transition probabilities are then given by Δ𝑓*_s,t_* for all sequences s,t:

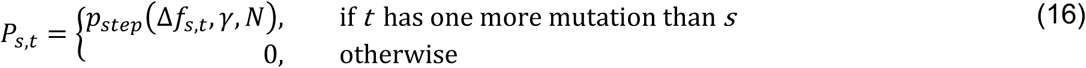

To compute the total probability for a specific antigen (all analyses used BA1 affinities), we compute the matrix product over all mutational steps *i* from the germline (*s_g_*) to the somatic (*s_s_*) sequence:

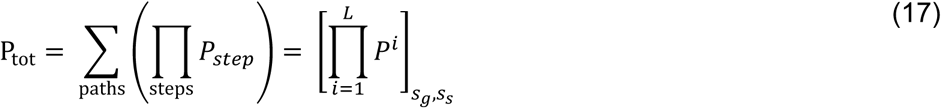

These probabilities are not normalized (*i.e.,* mutations are optional, and many paths will not reach the somatic state), and they encode the probability of reaching the somatic state. Because these are not true probabilities, they have arbitrary units and thus cannot be compared across models. However, *P_tot_* in the strong selection model can be interpreted as the number of uphill paths, and this number can be compared across selection models.

For all models, we perform 10 bootstraps, resampling each binding affinity, expression, and polyspecificity value from a normal distribution, recalculate *P_tot_* and average over 10 bootstrapped values to obtain the mean and SEM. Probabilities from the strong selection model are presentED **Fig. 2a,c**; probabilities from the weak selection model are presented in **Fig. 2b,d,e**.

To identify the most likely paths in each model, we use directed graphs where each sequence *s* is a node with directed edges toward all sequences *t* that can be reached in a single mutation, and edge weights are computed directly from the transition probability. We then use the Python package *networkx*^93^ to compute the most likely paths^94^. To compute the probability that a mutation at site *m* happened at a specific step *j*, we normalize the transition matrices so that each row sums to one and define:

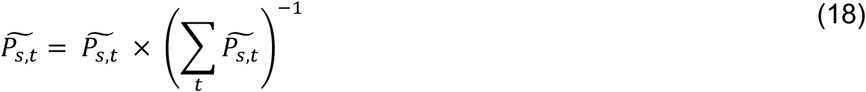

if 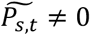 and 0 otherwise. The total relative probability for a site at a given mutational step is then:

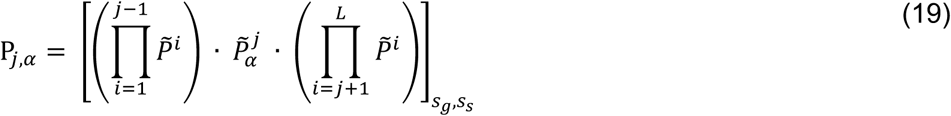

Because a sequence of L mutations starting from the germline can only lead to Omi32, 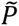 verifies 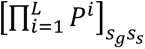 = 1. Thus, these probabilities are normalized over all sites and are summarized in **Figure 2d**.

Finally, we compute the total likelihood of each variant *s* containing *j* mutations by summing the probability of all paths containing that variant:

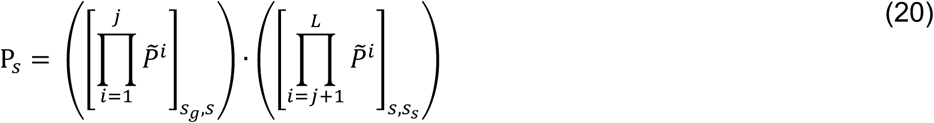

where the first term is the probability of reaching sequence *s* at mutational step *j* and the second term is the probability of reaching the somatic sequence after passing through sequence *s*. This probability is normalized to the probability in a neutral model (1/*n_j_*) such that variants with different numbers of mutations have comparable values. These probabilities are mapped onto force-directed layouts in **Fig. 2e**.

### Fab-spike structural characterization

#### Fab production and purification

Heavy and light chains were cloned into a human cell protein expression vector (**SI Files 6-9**). Plasmids encoding the heavy chain (57.1 µg) and light chain (142.9 µg) were co-transfected into HEK Expi293F cells (Thermo Fisher A14257; 200 µg total plasmid DNA per 600 million cells) according to the manufacturer’s instructions. IgG-containing media was harvested 6 days post-transfection and clarified by centrifugation at 300 × g for 10 min at 4°C, followed by a second spin at 4,000 × g for 15 min at 4°C. The supernatant was filtered through a 0.45 µm SFCA filter and diluted 1:1 with 1× PBS (Corning 21-040-CM).

Full-length IgG was purified using a 5 mL Cytiva Protein A HP column (Cytiva 17040301) on an ÄKTA go FPLC system according to the manufacturer’s instructions (binding buffer: 1× PBS; elution buffer: Pierce IgG Elution Buffer, Thermo Scientific 21004). Immediately following elution, IgG-containing fractions (identified by absorbance at 280 nm) were neutralized with 1:10 (v/v) 1 M Tris·HCl, pH 8.5 (from NAb™ Protein A Plus Spin Kit, Thermo Scientific 89948). Eluate fractions were pooled and concentrated to 5 mL using a 50 kDa molecular weight cutoff concentrator (Sigma-Aldrich UFC9050) by centrifugation at 4,800 × g at 4°C.

The concentrated IgG was desalted into papain sample buffer (20 mM sodium phosphate monobasic, 10 mM EDTA, pH 7.0 at 25°C) using PD-10 desalting columns (Cytiva 17085101). Desalted protein (∼5 mL) was diluted 1:1 with freshly prepared papain digestion buffer (20 mM cysteine-HCl in sample buffer, pH 7.0 at 25°C) and incubated with 4 mL immobilized papain (Thermo Scientific 20341) for 16 h at 37°C with end-over-end rotation.

Following papain proteolysis, the reaction pH was adjusted by addition of 5 mM Tris·HCl, pH 7.5, and the digest was reapplied to a Cytiva Protein A HP column. Flow-through and wash fractions containing Fab were pooled and concentrated using a 30 kDa molecular weight cutoff concentrator (Sigma-Aldrich UFC9030). Final purification was performed by size-exclusion chromatography on a Superose 6 Increase 10/300 GL column (Cytiva 29091596). Fractions containing Fab were pooled and concentrated to 2–5 mg/mL, as determined by absorbance at 280 nm using an extinction coefficient of 68,925 M⁻¹cm⁻¹ and the calculated Fab molecular weight of 47 kDa. Aliquots were stored at −80°C. Purity was assessed by SDS-PAGE followed by Coomassie staining.

#### Spike production and purification

Spike-6P (spike ectodomain with hexa pro stabilizing mutations and furin cleavage deleted^95^ - thrombin cleavage site - foldon - 6XHIs - avi-tag) in a mammalian protein expression plasmid (**SI Files 10-11**) was transfected into HEK Expi293F cells (ThermoFisher A14257; 200 ug plasmid per 600 million cells), per the manufacturer’s instructions. Spike-containing media was harvested 4 days post-transfection, spun at 3500 x g for 15 min at 4°C, filtered through a 0.22 micron SFCA filter, and purified by NiNTA chromatography, using a Cytiva His Trap FF column (Cytiva #17531901), per the manufacturer’s instructions. Eluted protein was then concentrated using a 50 kDa filter to < 1 mL (Millipore Sigma #UFC905008) before performing SEC in TBS. Purified Spike-6P was snap-frozen and thawed just before use.

#### Grid preparation and data collection for Spike+Fab+LC-Kappa VHH

Grids were prepared on UltrAuFoil 1/1 300 mesh holey gold grids (Electron Microscopy Sciences) and plunged on a Thermo Fisher Scientific Vitrobot Mk IV cryo-plunger. The final protein solution contained 4 μM BA1, 8 μM germline, 16 μM LC-Kappa VHH (Thermo Scientific 1033270500), 150 mM NaCl, 50 mM Tris pH 7.6. LC-Kappa was added to stabilize the antibody hinge region and reach a sufficient molecular weight for cryo-EM^96^. Prior to application on the grid, germline and LC-Kappa were mixed in a 1:2 molar ratio and incubated on ice for 30 min. BA1 spike was then added to a final molar ratio of 1:2:4 BA1:germline:LC-Kappa and incubated for an additional 30 min on ice. 3 μL of the final protein solution was applied to grids previously glow-discharged using a PELCO easiGlow.

Data were collected at NanoImaging Services (Woburn, MA) on a Thermo Fisher Glacios 200 kV electron microscope equipped with a Falcon 4 camera. 10,318 high-magnification images were collected 0.92 Å/pix, 19 frames, 24.85 e-/Å^2^ total dose, defocus range -1.0 to -2.5 μm.

The grid and collection parameters for BA4+Omi32+LC-Kappa, and BA4+germline+LC-Kappa are shown in **SI File 11 – Table 1**.

#### Grid preparation and data collection for Fab+LC-Kappa VHH

Grids were prepared on UltrAuFoil 0.6/1 300 mesh holey gold grids (Quantifoil) and plunged on a Thermo Fisher Scientific Vitrobot Mk IV cryo-plunger. The final protein solution contained 26 μM germline, 52 μM LC-Kappa, 150 mM NaCl, 50 mM Tris pH 7.6. Prior to application on the grid, germline and LC-Kappa were mixed in a 1:2 molar ratio and incubated on ice for 30 min. 3 μL of the final protein solution was applied to grids previously glow-discharged using a PELCO easiGlow.

Data were collected at NanoImaging Services (San Diego, CA) on a Thermo Fisher Titan Krios 300 kV electron microscope equipped with a Gatan K2 camera. 8,731 high-magnification images were collected 0.51 Å/pix, 51 frames, 49.91 e-/Å^2^ total dose, defocus range 0.4-1.8 μm.

The grid and collection parameters for Omi32+LC-Kappa are shown in **SI File 11 – Table 2**.

#### Data processing for Spike+Fab+LC-Kappa VHH

All Spike+Fab+LC-Kappa VHH datasets followed highly analogous processing workflows (**ED Fig. 9**), and the BA1+germline+LC-Kappa VHH workflow will be described here as a representative example. All data processing was performed in cryoSPARC 4^97^. Patch motion correction and contrast transfer function (CTF) estimation was performed in cryoSPARC live. ∼2.3 million particles were picked from a subset of 7,764 (micrographs with CTF fits lower than 5 Å were removed) images using cryoSPARC’s template picker and subjected to 2D classification, resulting in a set of ∼646k particles. These particles were subjected to ab initio classification, revealing classes with Fab occupancy ranging from zero to three per Spike trimer. The particles corresponding to the class with best density for Spike RBD+germline Fab were subjected to iterative heterogeneous refinement and non-uniform refinement, resulting in a set of ∼230k particles. The particles were then subjected to iterative rounds of 3D classification and 3D variability analysis, resulting in a best set of ∼16k particles, followed by global and local CTF refinement. The particles were symmetry-expanded to ∼48k particles (C3 symmetry) prior to being subjected to local refinement with a soft-padded mask around LC-Kappa, germline, and the RBD domain of BA1 (local refinement performed in C1). The final nominal resolution at FSC=0.143 is 3.2 Å. Following local refinement, model building was performed only for the RBD+Fab+LC-Kappa VHH. Data processing for BA4+Omi32+LC-Kappa and BA4+ germline+LC-kappa were performed analogously and resulted in final maps at 3.2 Å and 3.4 Å nominal resolution, respectively (**SI File 11 – Table 3**).

#### Data processing for Fab+LC-Kappa VHH

Both Fab+LC-Kappa VHH datasets followed highly analogous processing workflows (**ED Fig. 9**), and the germline+LC-Kappa VHH workflow will be described here as a representative example. All data processing was performed in cryoSPARC 4^97^. Patch motion correction and contrast transfer function (CTF) estimation was performed in cryoSPARC live. ∼2.7M particles were picked from a subset of 4,142 (micrographs with CTF fits lower than 5 Å were removed) images using a pre-trained Topaz model (as implemented in cryoSPARC) and subjected to 2D classification, resulting in a set of ∼211k particles. These particles were subjected to ab initio classification, heterogeneous refinement, and non-uniform refinement, resulting in a set of ∼93k particles, which were further subjected to global and local CTF refinements. A subset of ∼74k particles were selected and subjected to reference-based motion correction, followed by non-uniform refinement and local refinement with a soft-padded mask around germline and LC-Kappa. Particle classes with different loop conformations were not identified during processing. The final nominal resolution at FSC=0.143 is 3.1 Å. Data processing for Omi32+LC-Kappa was performed analogously, yielding a final map at a nominal resolution of 3.2 Å (**SI File 11 – Table 4**).

#### Model building and refinement

Model building and refinement for all samples were carried out analogously and initiated with published models for BA1 Spike RBD (PDB ID: 7ZFE^44^), BA4 Spike RBD (PDB ID: 7ZXU^45^), and Omi32 Fab (PDB ID: 7ZFE^44^). The starting model for germline was generated using AlphaFold^98^. The initial structures were placed into the sharpened density maps using the “fit to map” utility in ChimeraX^91^. Iterative rounds of model building and refinement were performed in PHENIX v.1.21.2^99^ and COOT v. 0.9.6 EL^100^. The final models were validated against the half-maps and its quality assessed by MolProbity^101^. For a full summary of refinement statistics, see **SI File 11 – Tables 3-4**.

### Biolayer interferometry affinity measurements

Biolayer interferometry measurements were made on an Octet RED384 (Sartorius). Biotinylated spike RBD was loaded onto Octet streptavidin (SA) biosensors (Sartorius #18-5019). All measurements were made in PBS supplemented with 1% BSA. Binding measurements were acquired as follows with shaking at 1000 rpm – sensor check: 60 s, baseline: 60 s, ligand loading: ∼60 s (until a density of ∼0.8 nm was acquired), second baseline: 60 s, association (600 s), and dissociation (600 s). Kinetics measurements for both Omi32 and germline Fabs were obtained at 26°C. Each Fab was assayed at 8 different spike RBD concentrations. Omi32 was assayed at 100 nM, 56 nM, 31 nM, 17 nM, 9.5 nM, 5.3 nM, 2.9 nM, and 0 nM for each spike RBD, and germline was assayed at 10 uM, 5.0 uM, 2.5 uM, 1.3 uM, 630 nM, 310 nM, 160 nM, and 0 nM. Equilibrium responses (*R*_eq_) were calculated by taking the final data point from the association phase at each spike RBD concentration, *c*, and fit to a 1:1 steady-state binding model to estimate 𝐾*_D_*:

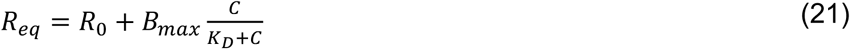

Where *B_max_* is the inferred maximum binding signal.

### Differential scanning fluorimetry stability measurements

Purified Fab samples were diluted to 2 mg/mL in PBS (Corning 21-040-CM), and concentrations were determined by absorbance at 280 nm using an extinction coefficient of 68,925 M⁻¹cm⁻¹ and the calculated Fab molecular weight of 47 kDa. Samples were then diluted to a final concentration of 1 mg/mL in PBS and mixed with 1× GloMelt dye (Biotium 33022-T) in a white 96-well plate (Roche 04729692001) to a total volume of 20 µL per well, in technical triplicates. All steps were performed on ice prior to loading the plate onto the instrument. Thermal melt experiments were conducted on a Roche LightCycler 480 qPCR machine. Plates were held at 25°C for 30s and scanned from 25°C to 99°C at a rate of 0.11°C/s, with detection set to SYBR Green I / HRM dye mode (excitation 465 nm, emission 510 nm). Raw fluorescence values were smoothed using the savgol_filter function in SciPy, and the first derivative (dF/dT) was calculated. The apparent melting temperature (T_m_^App^) was defined as the temperature corresponding to the maximum of the dF/dT curve, which was normalized to the germline sequence and is reported as the mean difference of three independent measurements performed on different days.

## Data availability

Data and code for this work are available at Github (https://github.com/Ctharp17/Omi32_BioPhy-Seq.git). Antibody affinity, polyreactivity, and expression data are also available in an interactive data browser at https://amphilli.github.io/Omi32_browser_git/. FASTQ files will be deposited in the NCBI BioProject database upon publication. Atomic models and EM maps were deposited to the Protein Data Bank (PDB) and Electron Microscopy Data Bank (EMDB), respectively. PDB and EMDB codes for the complexes reported here are: germline+LCKappa: PDB ID 11OU, EMD-75893; Omi32+LCKappa: PDB ID 11OR, EMD-75892; germline+BA1+LCKappa: PDB ID 11OQ, EMD-75891; germline+BA4+LCKappa: PDB ID 11OO; EMD-75889; Omi32+BA4+LCKappa: PDB ID 11OL, EMD-75887.

## Supplemental Files

Supplemental File 1. Plasmid map for integrating germline antibody into *attb* landing pad

Supplemental File 2. Plasmid map for integrating Omi32 antibody into *attb* landing pad

Supplemental File 3. Primers for combinatorial library generation

Supplemental File 4. Primers for Illumina sequencing library preparation

Supplemental File 5. Schematic of fluorescence-activated cell sorting for BioPhy-Seq measurements

Supplemental File 6. Plasmid map for recombinant expression of germline antibody (light chain)

Supplemental File 7. Plasmid map for recombinant expression of germline antibody (heavy chain)

Supplemental File 8. Plasmid map for recombinant expression of Omi32 antibody (light chain)

Supplemental File 9. Plasmid map for recombinant expression of Omi32 antibody (heavy chain)

Supplemental File 10. Plasmid map for recombinant expression of BA1 spike trimer

Supplemental File 11. Plasmid map for recombinant expression of BA4 spike trimer

Supplemental File 12. Supplemental Tables 1-4 (cryo-electron microscopy imaging conditions and refinement statistics)

Supplemental File 13. Video of antibody preconfiguration and antigen binding

---

**Extended Data Fig. 1.**
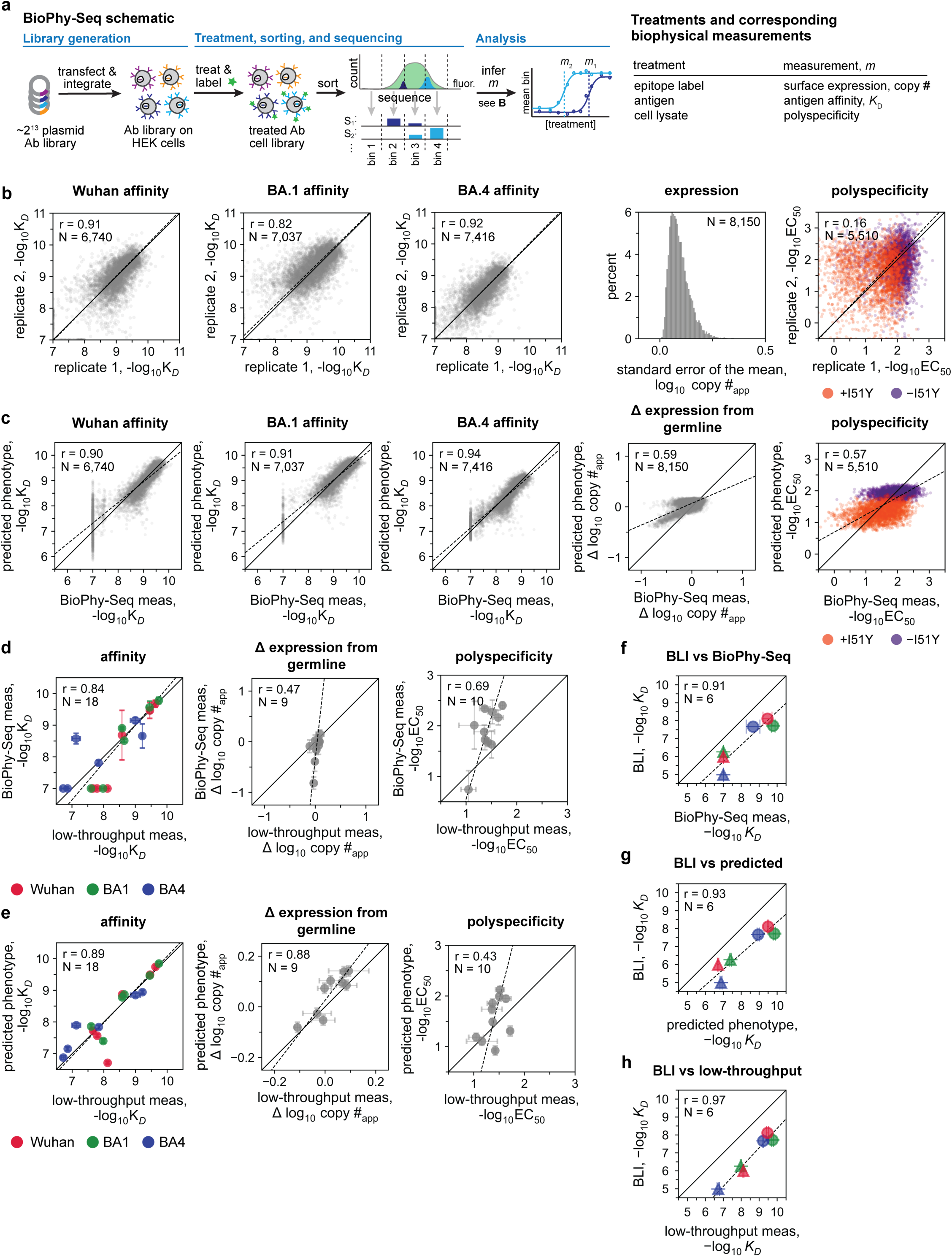
| BioPhy-Seq enables high-resolution measurements of antibody expression, antigen affinity, and polyspecificity. **a,** BioPhy-Seq schematic. Antibody (Ab) sequences are cloned into an epitope-tagged DNA-barcoded IgG backbone. Plasmid libraries are transfected and integrated into HEK cells with a genomic landing pad (only one sequence can integrate per cell). Cell libraries are labeled with fluorophore-conjugated treatment, flow sorted into bins based on fluorescence, and DNA barcodes are sequenced. Biophysical measurements (*m*) are inferred by computing the mean bin for each sequence in each treatment concentration. Measurements and treatments for this study are shown on the right. **b,** Correspondence between BioPhy-Seq measurements across biological replicates for Wuhan affinity, BA1 affinity, BA4 affinity (two replicates), expression (six replicates), and polyspecificity (two replicates). Due to experimental noise, polyspecificity measurements were less reproducible across BioPhy-Seq biological replicates. Still, given the large number of measurements (N=5,510, r=0.16), a second-order linear model captured significant first-order and pairwise interactions (r = 0.57; see **c** and Figure 3), and low-throughput clonal measurements recapitulated the BioPhy-Seq mean across replicates (r = 0.69; see **d**). To illustrate the large first-order effect of I51Y, **b-c** are colored by its identity across the library (I51 purple; Y51 orange). **c,** Correlation between BioPhy-Seq measurements and predicted values from linear interaction models for Wuhan affinity, BA1 affinity, BA4 affinity, difference in expression from germline, and polyspecificity; points indicate mean between biological replicates. **d,** Correlation between BioPhy-Seq and low-throughput flow cytometry measurements for affinity to Wuhan, BA1, and BA4, difference in expression from germline, and polyspecificity. Ten clonal cell lines expressing selected antibody intermediates (including Omi32 and germline) were used for low-throughput measurements; points and error bars indicate the mean and SEM between biological replicates, respectively (affinity SEM are generally smaller than the points). **e,** Correlation between predicted values from linear interaction models and low-throughput flow cytometry measurements of affinity (to Wuhan, BA1, and BA4), difference in expression from germline, and polyspecificity. Ten clonal cell lines expressing selected antibody intermediates (including Omi32 and germline) were used for low-throughput measurements; points and error bars indicate the mean and SEM between biological replicates, respectively (affinity SEM are generally smaller than the points). **f,** Correlation between BioPhy-Seq measurements and biolayer interferometry (BLI) for Omi32 (circles) and germline (triangles) affinities (Wuhan (red), BA1 (green) and BA4 (blue)); points and error bars indicate the mean and SEM for biological replicates, respectively (SEM are generally smaller than the points). **g,** Correlation between predicted values from linear interaction models and biolayer interferometry (BLI) for Omi32 (circles) and germline (triangles) affinities (Wuhan, BA1 and BA4); points and error bars indicate the mean and SEM for biological replicates, respectively (SEM are generally smaller than the points). **h,** Correlation between low-throughput flow cytometry measurements and biolayer interferometry (BLI) for Omi32 (circles) and germline (triangles) affinities (Wuhan, BA1 and BA4); points and error bars indicate the mean and SEM for biological replicates, respectively (SEM are generally smaller than the points). **For correlation plots in b-h, Pearson’s correlation coefficient (r), solid identity (1:1) line, and dashed least-squares regression line are shown on plot.

**Extended Data Fig. 2.**
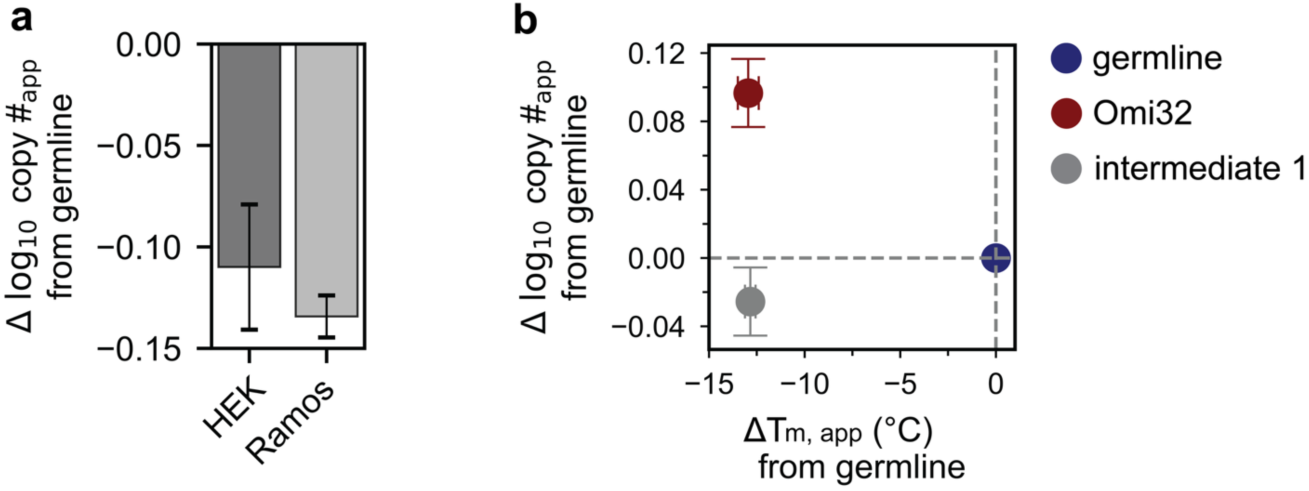
| Differences in expression measurements made with BioPhy-Seq correspond to expression differences in Ramos B-cells but not necessarily differences in thermal stability. **a,** Differences in expression between an influenza bnAb (CH65)^66^ and its corresponding germline sequence (UCA860) on HEK cells and Ramos B-cells. Bars and error bars indicate the mean and SEM between biological replicates (N=7), respectively. **b,** Correspondence between differences in HEK cell-surface expression and thermal stability (compared to germline) for Omi32 and a selected intermediate. Points and error bars indicate the mean and SEM between biological replicates (N=3), respectively (germline is included as a reference and therefore has no error bars)

**Extended Data Fig. 3.**
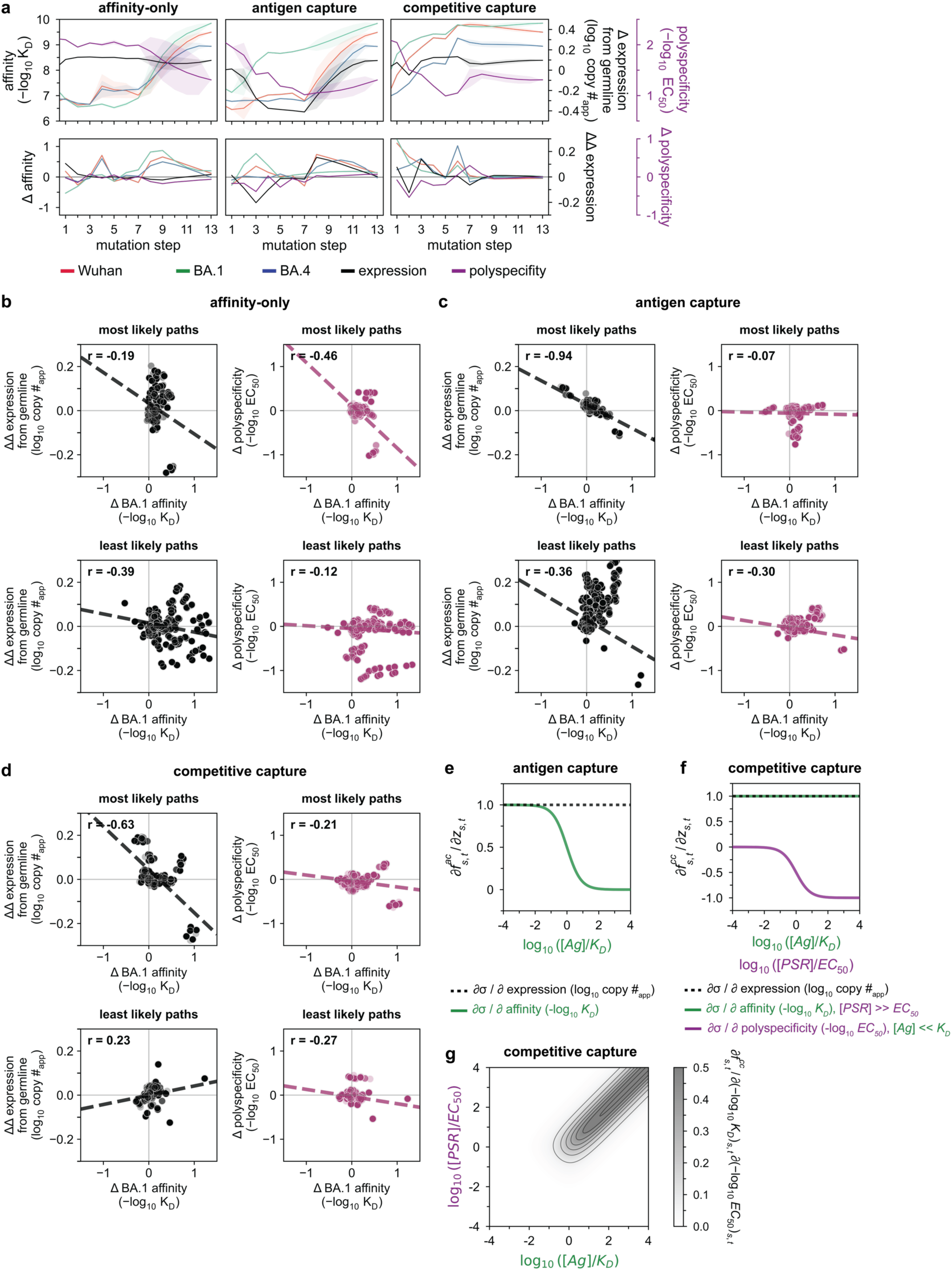
| Phenotype trajectories for least likely paths, correlations between mutation effects across paths, and selection of pathway model parameters. **a,** Top: mean phenotype at each mutational step for the least likely 1,000 paths under the three selection models (all models implement weak selection pressure where neutral and deleterious mutations are disfavored but allowed – see Methods for details). Shaded area represents one standard deviation from the mean. Bottom: mean change in phenotype at each step (bottom). **b,** Correlation between changes in affinity and expression (left) and affinity and polyspecificity (right) at each step for the most likely (top) and least likely (bottom) 1,000 paths for the affinity-only model (N=13,000). **c,** Correlation between changes in affinity and expression (left) and affinity and polyspecificity (right) at each step for the most likely (top) and least likely (bottom) 1,000 paths for the antigen capture model (N=13,000). **d,** Correlation between changes in affinity and expression (left) and affinity and polyspecificity (right) at each step for the most likely (top) and least likely (bottom) 1,000 paths for the competitive capture model (N=13,000). **e,** Graphical representation for the impact of antigen concentration ([𝐴𝑔]) on the selection coefficient (σ) for the antigen capture model (see Methods): Partial derivative of antigen capture fitness (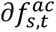) with respect to relevant antibody properties (∂𝑧*_s,t_)* at varying [𝐴𝑔]/𝐾*_D_* ratios indicates that the [𝐴𝑔] ≪ 𝐾*_D_* regime used for this study reduces to 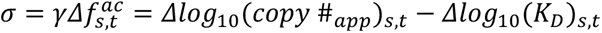 (*i.e.,* that improvements in antigen capture are dependent on improvements in copy number and binding affinity). **f,** Graphical representation for the impact of antigen concentration ([𝐴𝑔]) and polyspecificity reagent concentration ([*PSR*]) on the selection coefficient (σ) for the competitive capture model (see Methods): Partial derivative of competitive capture fitness (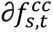) with respect to relevant antibody properties (∂𝑧*_s,t_*). Varying [𝐴𝑔]/𝐾*_D_* ratios at fixed [𝑃𝑆𝑅] ≫ 𝐸𝐶_50_ for affinity and varying [𝑃𝑆𝑅]/𝐸𝐶_50_ ratios at fixed [𝐴𝑔] ≪ 𝐾*_D_* (expression reduces to 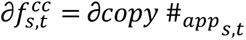 for both ratios) indicates that the [𝐴𝑔] ≪ 𝐾*_D_*, [𝑃𝑆𝑅] ≫ 𝐸𝐶_50_ regime used for this study reduces to 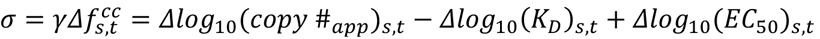 (*i.e.,* that improvements in competitive capture are dependent on copy number, binding affinity, and polyspecificity). **g,** Contour plot showing non-additive effects (mixed partial derivative) on competitive capture fitness (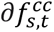) with respect to ∂𝑙𝑜𝑔_10_(𝐾*_D_*)*_s,t_* and ∂𝑙𝑜𝑔_10_(𝐸𝐶_50_)*_s,t_* at different combinations of [𝐴𝑔]/𝐾*_D_* and [𝑃𝑆𝑅]/𝐸𝐶_50_ ratios, confirming that the [𝐴𝑔] ≪ 𝐾*_D_*, [𝑃𝑆𝑅] ≫ 𝐸𝐶_50_ regime used for this study (upper-left quadrant) reduces to 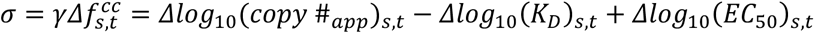 (*i.e.,* that improvements in competitive capture are dependent on copy number, binding affinity, and polyspecificity). **For b-d, all models implement weak selection pressure where neutral and deleterious mutations are disfavored but allowed – see Methods for details. Dashed line shows least-squares regression line.

**Extended Data Fig. 4.**
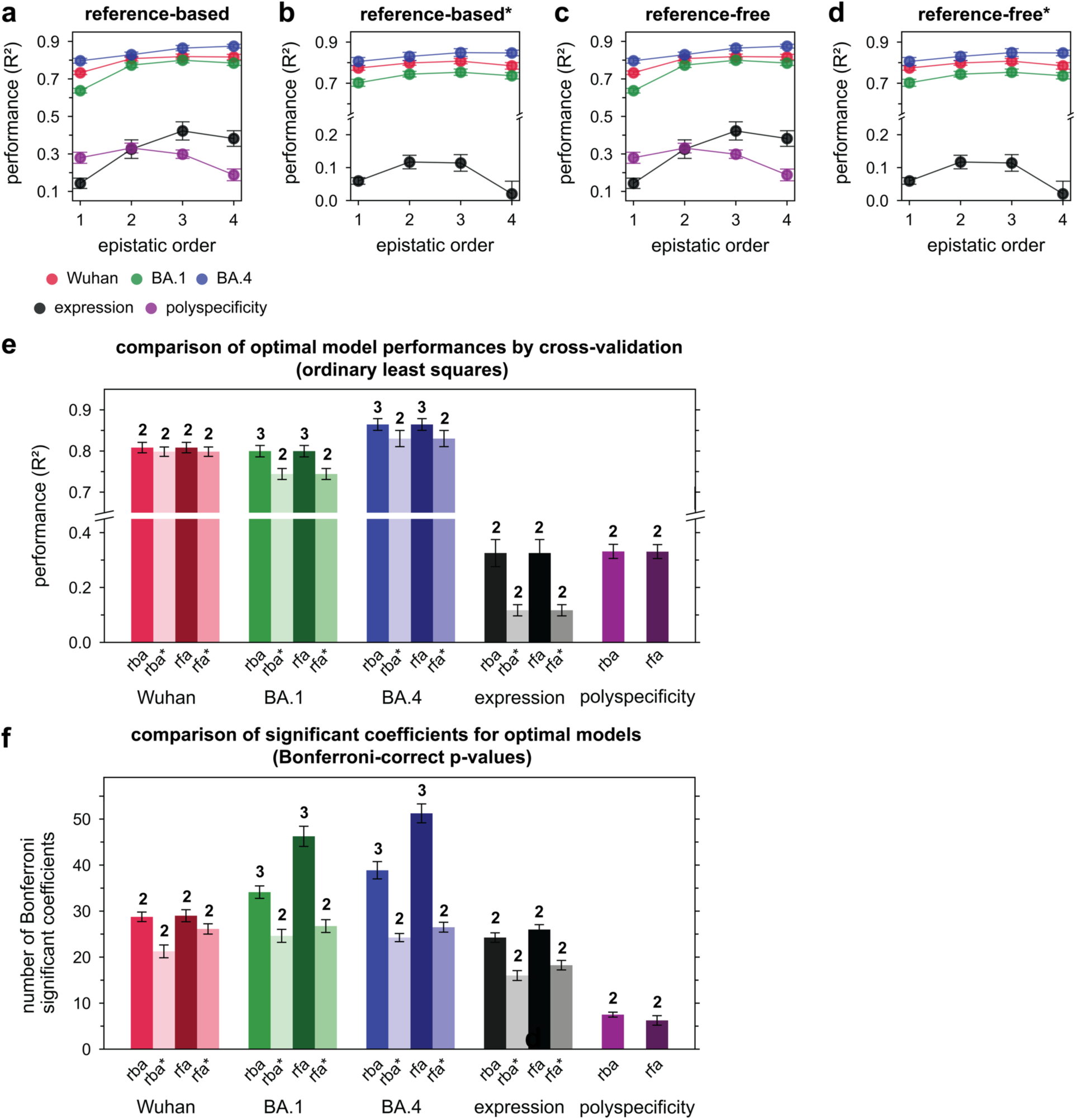
| Cross-validation results for reference-free and reference-based linear interaction models, with and without initial nonlinear transformation. **a,** Cross-validation performance of reference-based linear interaction models (see Methods). **b,** Cross-validation performance of reference-based linear interaction models following a nonlinear transformation (indicated by *) for each dataset (see Methods). **c,** Cross-validation performance of reference-free linear interaction models following a nonlinear transformation (indicated by *) for each dataset (see Methods). **d,** Cross-validation performance of reference-free linear interaction models following a nonlinear transformation (indicated by *) for each dataset (see Methods). **e,** Comparison of optimal model performance for each phenotype and each linear interaction model, with and without initial nonlinear transformation. Bars and error bars indicate the mean and SEM prediction performance (*R^2^*) across eight folds of ordinary least-squares cross-validation. Numbers on top of each bar indicate the maximum epistatic order inferred for each optimal model. Nonlinear transformations of the polyspecificity dataset were not used due to extreme bimodality in the data; reference-based analysis (rba), reference-based analysis with initial nonlinear transformation (rba*), reference-free analysis (rfa), reference-free analysis with initial nonlinear transformation (rfa*). **f,** Comparison of total Bonferroni-corrected significant coefficients (see Methods) inferred from optimal models for each phenotype and each linear interaction model, with and without initial nonlinear transformation. Bars and error bars indicate the mean and SEM of total Bonferroni-corrected significant coefficients across eight folds of ordinary least-squares cross-validation. Numbers on top of each bar indicate the maximum epistatic order inferred for each optimal model. Nonlinear transformations of the polyspecificity dataset were not used due to extreme bimodality in the data; reference-based analysis (rba), reference-based analysis with initial nonlinear transformation (rba*), reference-free analysis (rfa), reference-free analysis with initial nonlinear transformation (rfa*). **For a-d, points and error bars indicate the mean and SEM prediction performance (*R^2^*) across eight folds of ordinary least-squares cross-validation.

**Extended Data Fig. 5.**
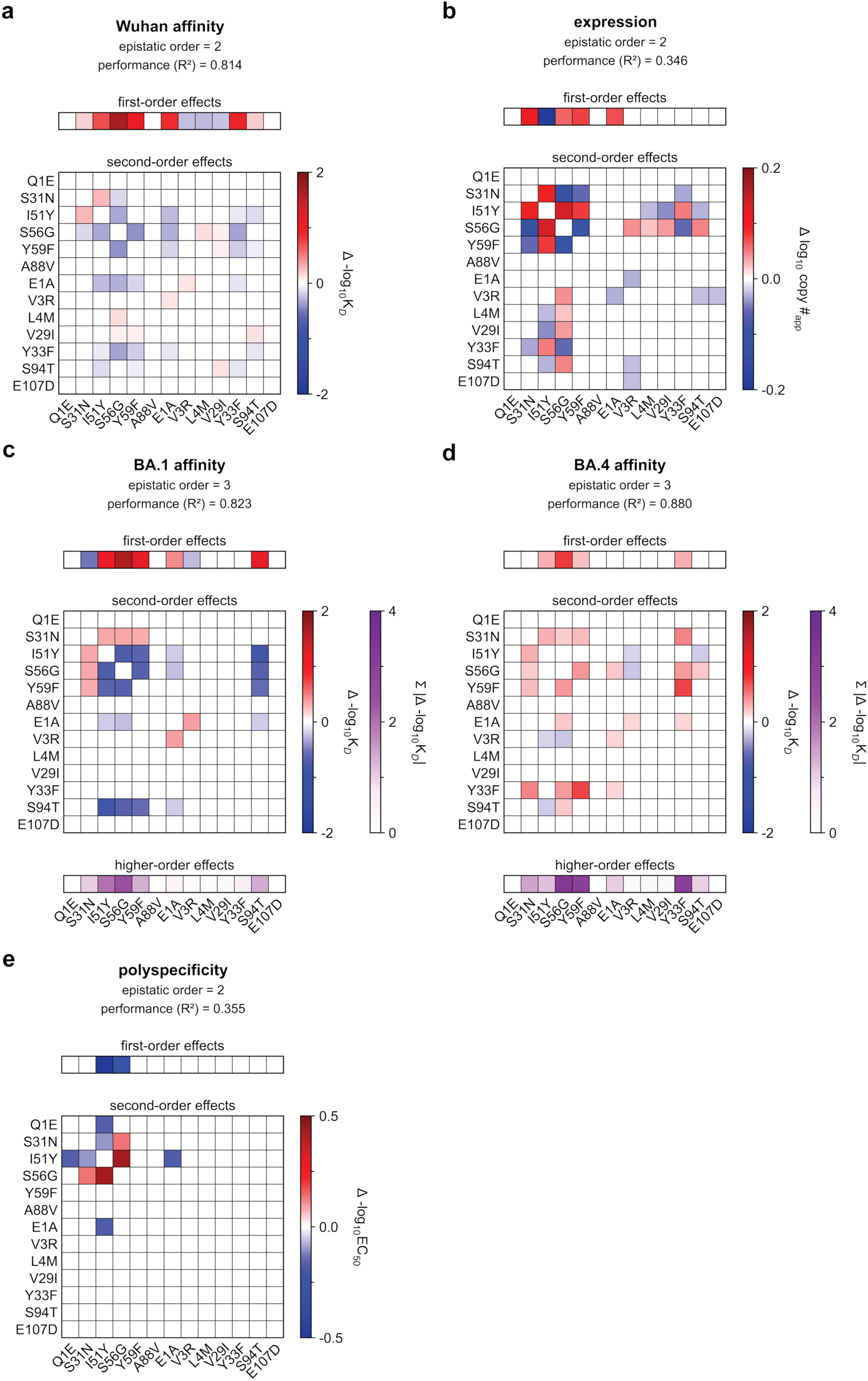
| Epistatic coefficients for the optimal epistatic model for each phenotype using reference-based linear interaction models. **a,** First and second-order interaction strengths for the optimal (epistatic order = 2) reference-based linear interaction model for the Wuhan affinity dataset. **b,** First and second-order interaction strengths for the optimal (epistatic order = 2) reference-based linear interaction model for the expression dataset. **c,** First, second, and higher-order interaction strengths for the optimal (epistatic order = 3) reference-based linear interaction model for the BA1 affinity dataset. **d,** First, second, and higher-order interaction strengths for the optimal (epistatic order = 3) reference-based linear interaction model for the BA4 affinity dataset. **e,** First and second-order interaction strengths for the optimal (epistatic order = 2) reference-based linear interaction model for the polyspecificity dataset. **For a-e, Omi32 mutations are labeled on the axes. First and second-order coefficient cells are colored by the blue-white-red color bar only if the epistatic coefficient is significant (Bonferroni-corrected p-value). The higher-order effect cells are colored according to the white-purple color bar and indicate the sum of absolute Bonferroni-corrected significant coefficients including each Omi32 mutation. *R^2^*: model predictive performance for the entire dataset.

**Extended Data Fig. 6.**
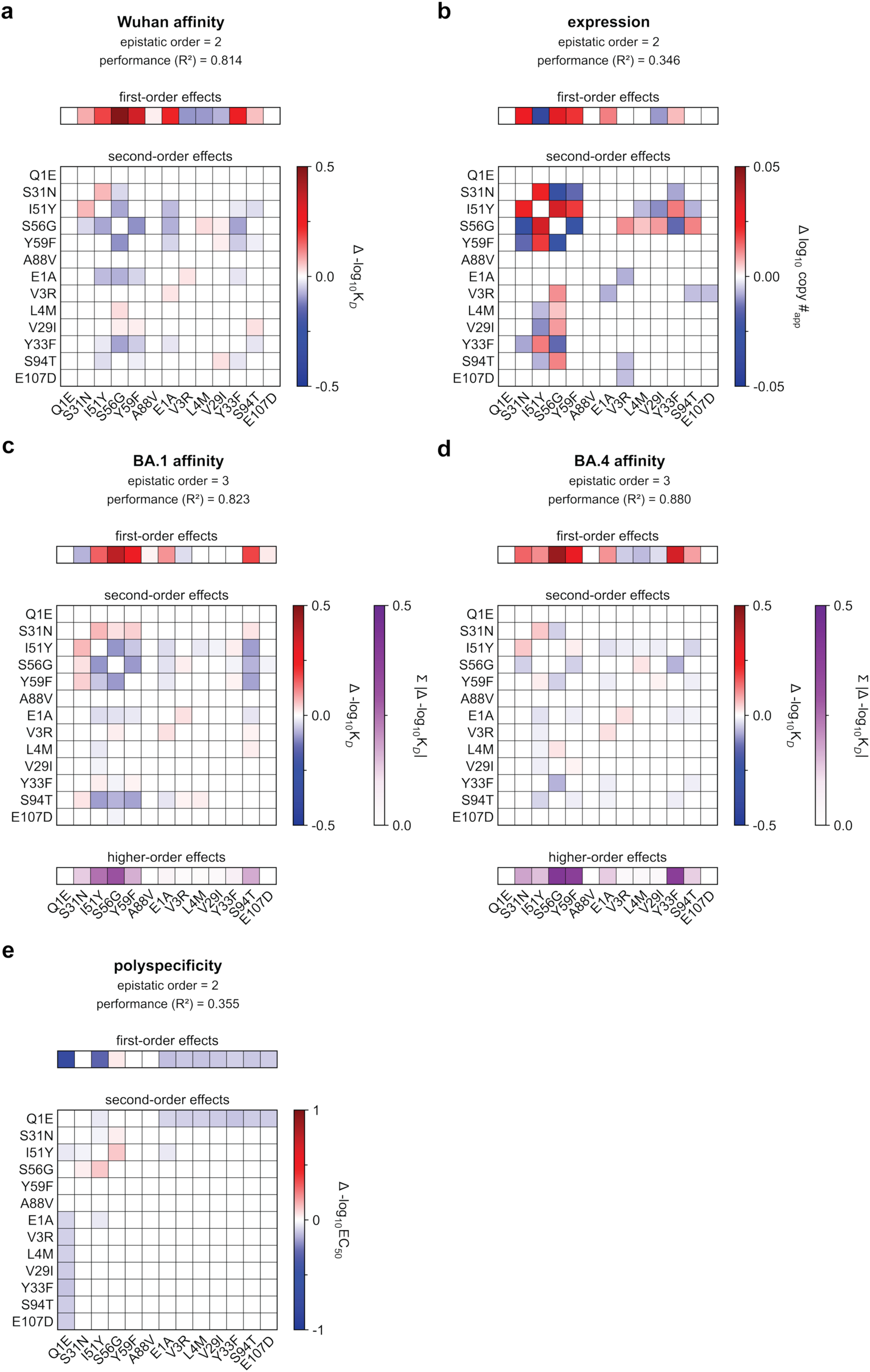
| Epistatic effects for the optimal epistatic model for each phenotype using reference-free linear interaction model. **a,** First and second-order interaction strengths for the optimal (epistatic order = 2) reference-free linear interaction model for the Wuhan affinity dataset. **b,** First and second-order interaction strengths for the optimal (epistatic order = 2) reference-free linear interaction model for the expression dataset. **c,** First, second, and higher-order interaction strengths for the optimal (epistatic order = 3) reference-free linear interaction model for the BA1 affinity dataset. **d,** First, second, and higher-order interaction strengths for the optimal (epistatic order = 3) reference-free linear interaction model for the BA4 affinity dataset. **e,** First and second-order interaction strengths for the optimal (epistatic order = 2) reference-free linear interaction model for the polyspecificity dataset. **For a-e, Omi32 mutations are labeled on the axes. First and second-order coefficient cells are colored by the blue-white-red color bar only if the epistatic coefficient is significant (Bonferroni-corrected p-value). The higher-order effect cells are colored according to the white-purple color bar and indicate the sum of absolute Bonferroni-corrected significant coefficients including each Omi32 mutation. *R^2^*: model predictive performance for the entire dataset.

**Extended Data Fig. 7.**
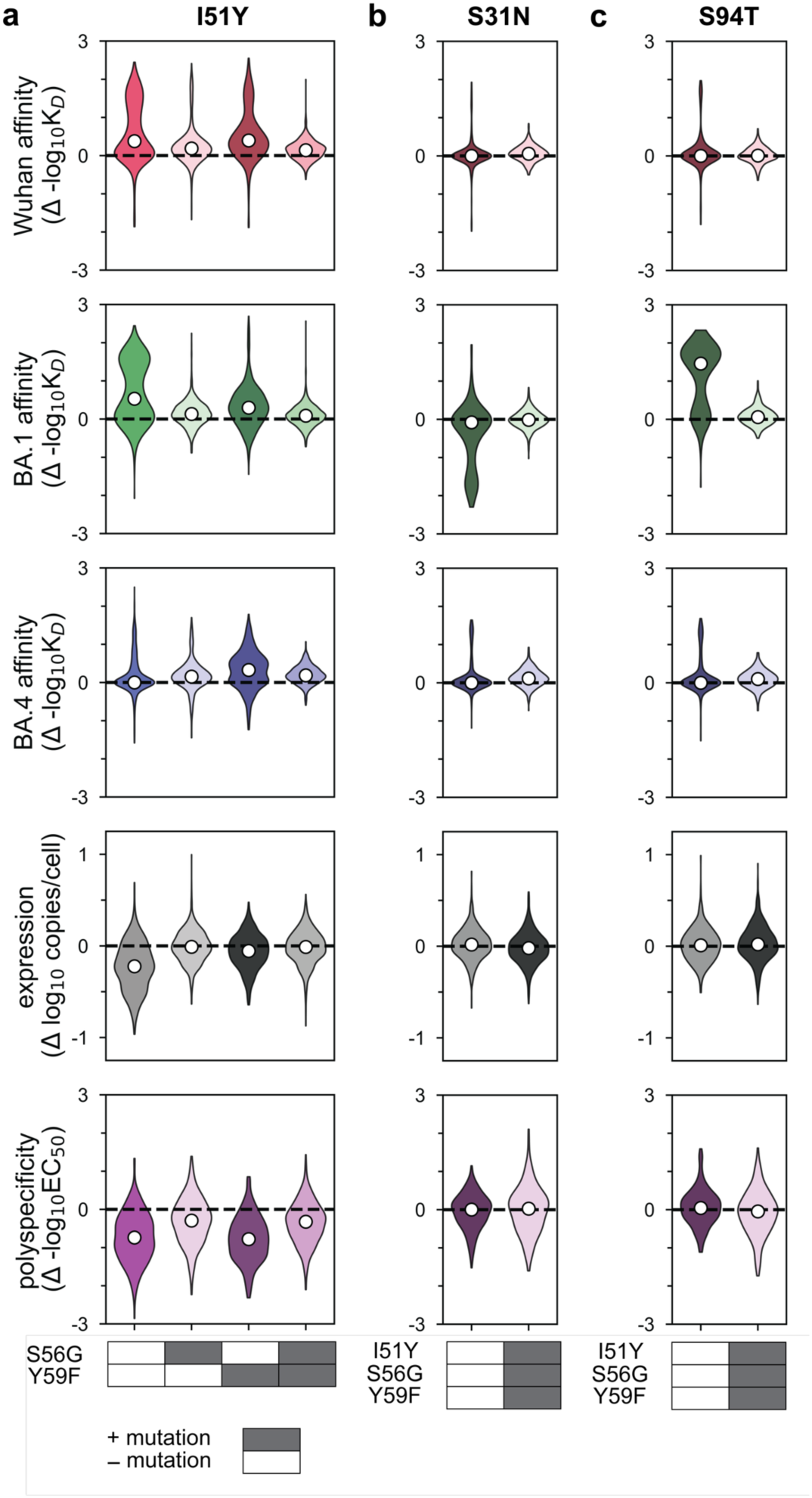
| Phenotypic effects of I51Y, S31N, and S94T with and without heavy-chain HCDR2 mutations. **a,** Effect of I51Y on experimental measurements of Wuhan affinity, BA1 affinity, BA4 affinity, expression, and polyspecificity for sequences with and without S56G and Y59F (N∼1024 genotypes per violin). **b,** Effect of S31N on experimental measurements of Wuhan affinity, BA1 affinity, BA4 affinity, expression, and polyspecificity for sequences with and without I51Y, S56G, and Y59F (N∼512 genotypes per violin). **c,** Effect of S94T on experimental measurements of Wuhan affinity, BA1 affinity, BA4 affinity, expression, and polyspecificity for sequences with and without and I51Y, S56G, Y59F (N∼512 genotypes per violin). **For a-c, black line indicates zero effect. White dots indicate the median for each violin. Boxes underneath the plots indicate the presence (gray) or absence (white) of the specified mutation.

**Extended Data Fig. 8.**
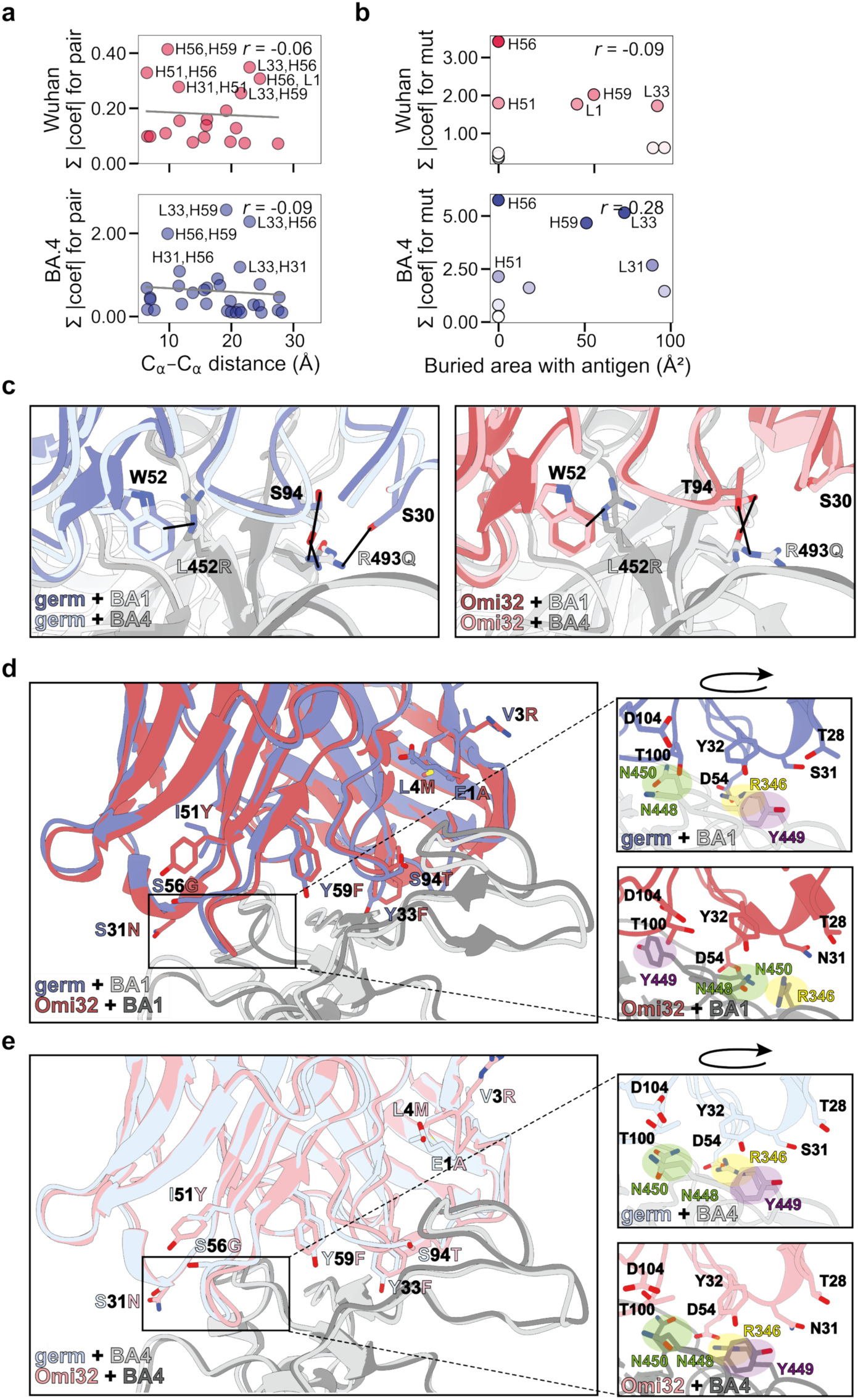
| Structural analysis of antibody-antigen interactions. **a,** Relationship between epistatic effects and distance between pairs of residues, computed from Omi32-BA1 structure (PDB ID 7ZFE)^44^, as in Fig. 4a. Pearson r shown on plot. **b,** Relationship between mutational effect size and buried area with the antigen. Some large-effect mutations make no contact with antigen, computed from Omi32-BA1 structure (PDB ID 7ZFE)^44^, as in Fig. 4b. Pearson r shown on plot. **c**, Structures of germline (left) and Omi32 (right) bound to BA1 and BA4. L452R and R493Q mutations that differentiate BA1 from BA4 are labeled; antibody residues that interact with these sites are also labeled, and those interactions are denoted with black lines. Global alignment was performed on heavy and light chains: germline+BA1 vs germline+BA4 RMSD (0.659 Å), Omi32+BA1 vs Omi32+BA4 RMSD (0.737 Å). **d**, Structures of germline and Omi32 bound to BA1. Alignment in the left panel shows that the overall conformations are similar; interface mutations in the antibodies are labeled. Zoomed view in the right panels shows that BA1 residues change orientation and interact with different residues in germline and Omi32, but the types of interactions with antibody residues are similar. Global alignment was performed on heavy and light chains, with an RMSD of 0.830 Å. **e**, Structures of germline and Omi32 bound to BA4. Alignment in the left panel shows that the overall conformations are similar; interface mutations in the antibodies are labeled. Zoomed view in the right panels shows that BA4 residues do not change orientation. Global alignment was performed on heavy and light chains, with an RMSD of 0.606 Å. *In c-e, all structures were solved by cryo-EM, except for the existing Omi32-BA1 structure solved by X-ray crystallography (PDB ID 7ZFE)^44^. Structures for germline, Omi32, germline+BA4, Omi32+BA4, and germline+BA1 were determined to 3.1 Å, 3.2 Å, 3.4 Å, 3.2 Å, 3.2 Å nominal resolution, respectively.

**Extended Data Fig. 9.**
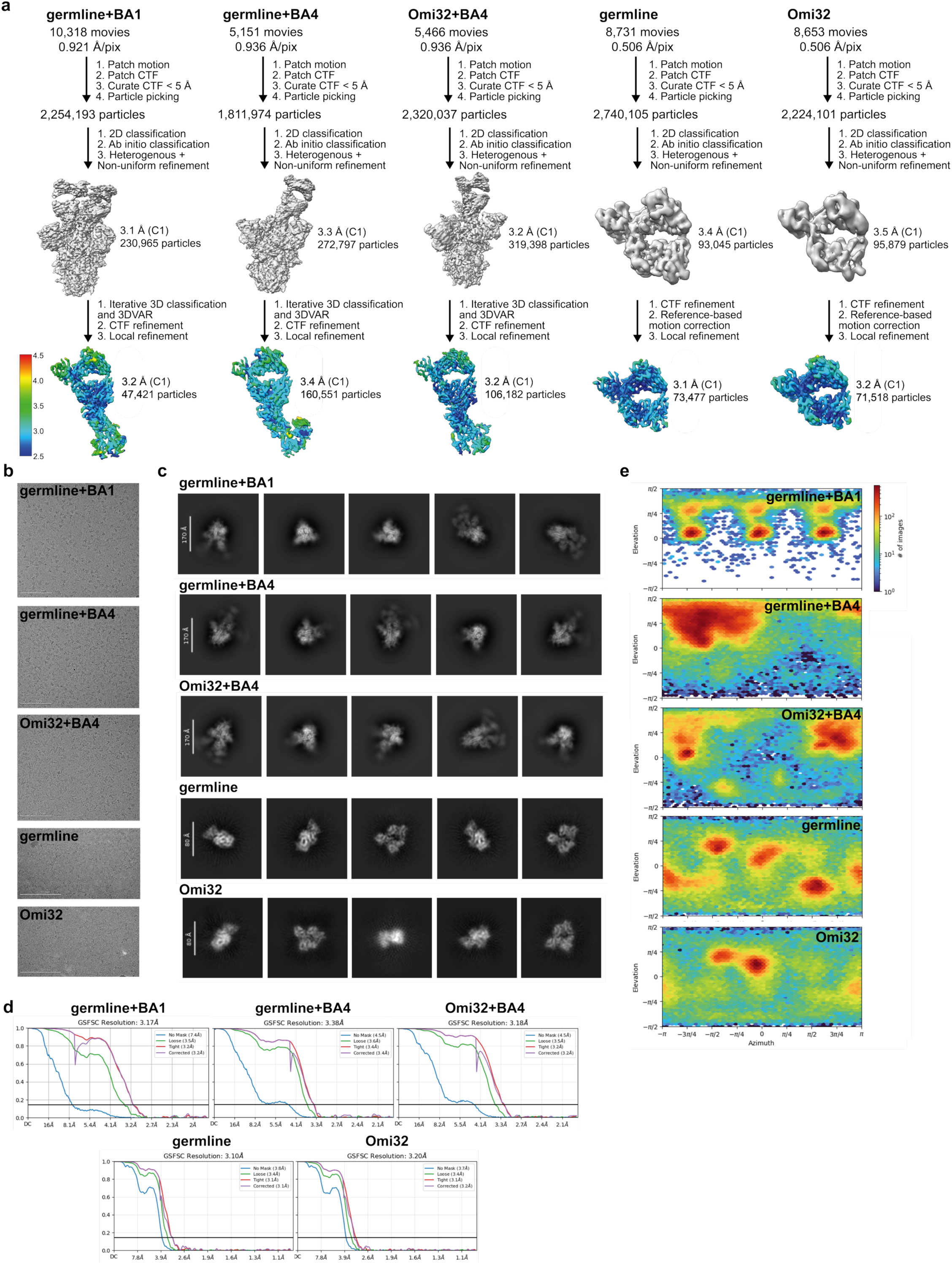
| Summary of cryo-EM data processing. **a,** Processing workflows **b**, Representative micrographs **c**, Representative 2D classes **d**, FSC curves for final maps (calculated by cryoSPARC)^97^ **e**, Viewing distributions for final maps (calculated by cryoSPARC)^97^ **\*\***All structures also contain the LC-Kappa VHH.

**Extended Data Fig. 10.**
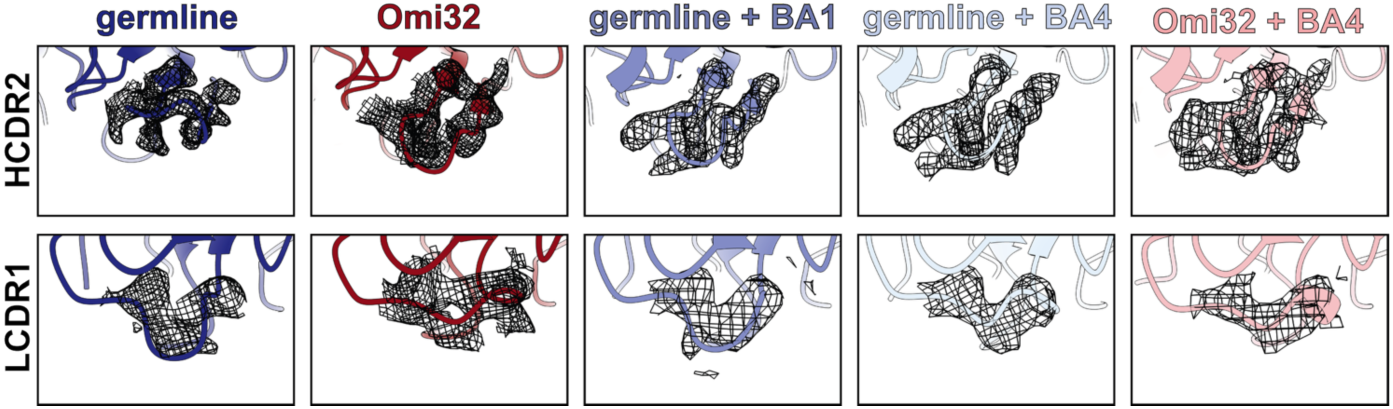
| HCDR2 and LCDR1 loop conformations for germline and Omi32, unbound or bound to BA1/BA4. Top: HCDR2 loop adopts a “down” conformation in Omi32, germline+BA1, germline+BA4, and Omi32+BA4, and adopts an “up” conformation in germline. Density around respective HCDR2 loops (residues 51-58 in the heavy chain) is shown in mesh. Bottom: LCDR1 loop adopts a “down” conformation in germline and germline+BA1, and adopts an “up” conformation in Omi32, germline+BA4, and Omi32+BA4. Density around respective LCDR1 loops (residues 29-32 in the light chain) is shown in mesh.

## References

1 Rotem, A. et al. Evolution on the biophysical fitness landscape of an RNA virus. Mol. Biol. Evol. 35, 2390, (2018)

2 Gong, L. I. et al. Stability-mediated epistasis constrains the evolution of an influenza protein. eLife 2, e00631, (2013)

3 Tokuriki, N. et al. Stability effects of mutations and protein evolvability. Curr. Opin. Struc. Biol. 19, 596, (2009)

4 Faure, A. J. et al. Mapping the energetic and allosteric landscapes of protein binding domains. Nature 604, 175, (2022)

5 Ghose, D. A., et al. Marginal specificity in protein interactions constrains evolution of a paralogous family. Proc. Nat. Acad. Sci. USA 120, e2221163120, (2023)

6 Muir, D. F. et al. Evolutionary-scale enzymology enables exploration of a rugged catalytic landscape. Science 388, eadu1058, (2025)

7 Burnett, D. L. et al. Germinal center antibody mutation trajectories are determined by rapid self/foreign discrimination. Science 360, 223, (2018)

8 Shehata, L. et al. Affinity maturation enhances antibody specificity but compromises conformational stability. Cell Rep. 28, 3300, (2019)

9 Victora, G. D. et al. Germinal centers. Annu. Rev. Immunol. 40, 413, (2022)

10 Horns, F., et al. Signatures of selection in the human antibody repertoire: Selective sweeps, competing subclones, and neutral drift. Proc. Nat. Acad. Sci. USA 116, 1261, (2019)

11 DeWitt, W. S., et al. Replaying germinal center evolution on a quantified affinity landscape. bioRxiv, (2025)

12 Batista, F. D. et al. Affinity dependence of the B cell response to antigen: a threshold, a ceiling, and the importance of off-rate. Immunity 8, 751, (1998)

13 Corti, D. et al. Broadly neutralizing antiviral antibodies. Annu. Rev. Immunol. 31, 705, (2013)

14 Kreer, C., et al. Probabilities of developing HIV-1 bNAb sequence features in uninfected and chronically infected individuals. Nat. Comm. 14, 7137, (2023)

15 Fu, Y., et al. A broadly neutralizing anti-influenza antibody reveals ongoing capacity of haemagglutinin-specific memory B cells to evolve. Nat. Comm. 7, 12780, (2016)

16 Chen, Y. et al. Broadly neutralizing antibodies to SARS-CoV-2 and other human coronaviruses. Nat. Rev. Immunol. 23, 189, (2023)

17 Cao, Y. et al. BA. 2.12. 1, BA. 4 and BA. 5 escape antibodies elicited by Omicron infection. Nature 608, 593, (2022)

18 Cohen, A. A. et al. Mosaic RBD nanoparticles protect against challenge by diverse sarbecoviruses in animal models. Science 377, eabq0839, (2022)

19 Sprenger, K. G., et al. Optimizing immunization protocols to elicit broadly neutralizing antibodies. Proc. Nat. Acad. Sci. USA 117, 20077, (2020)

20 Wang, S. et al. Manipulating the selection forces during affinity maturation to generate cross-reactive HIV antibodies. Cell 160, 785, (2015)

21 Julian, M. C. et al. Efficient affinity maturation of antibody variable domains requires co-selection of compensatory mutations to maintain thermodynamic stability. Sci. Rep. 7, 45259, (2017)

22 König, J., et al. Association-induced folding governs surrogate light chain and pre-B cell receptor core assembly. Nat. Comm. 17, 1202, (2026)

23 Feige, M. J. et al. How antibodies fold. Trends Biochem. Sci. 35, 189, (2010)

24 Xu, Y. et al. Addressing polyspecificity of antibodies selected from an in vitro yeast presentation system: a FACS-based, high-throughput selection and analytical tool. Protein Eng. Des. Sel. 26, 663, (2013)

25 Goodnow, C. et al. in Cold Spring Harbor symposia on quantitative biology. 907 (CSHL Press).

26 Kelly, R. L. et al. Nonspecificity in a nonimmune human scFv repertoire. MAbs 9, 1029, (2017)

27 Bajic, G. et al. Autoreactivity profiles of influenza hemagglutinin broadly neutralizing antibodies. Sci. Rep. 9, 3492, (2019)

28 Guthmiller, J. J. et al. Polyreactive broadly neutralizing B cells are selected to provide defense against pandemic threat influenza viruses. Immunity 53, 1230, (2020)

29 Tan, C. et al. Nur77 links chronic antigen stimulation to B cell tolerance by restricting the survival of self-reactive B cells in the periphery. J. Immunol. 202, 2907, (2019)

30 Mouquet, H. et al. Polyreactive antibodies in adaptive immune responses to viruses. Cell Mol. Life Sci. 69, 1435, (2012)

31 Boder, E. T. et al. Yeast surface display for screening combinatorial polypeptide libraries. Nat. Biotech. 15, 553, (1997)

32 Smith, G. P. Filamentous fusion phage: novel expression vectors that display cloned antigens on the virion surface. Science 228, 1315, (1985)

33 Adams, R. M. et al. Measuring the sequence-affinity landscape of antibodies with massively parallel titration curves. eLife 5, e23156, (2016)

34 Schaefer, J. V. et al. Transfer of engineered biophysical properties between different antibody formats and expression systems. Protein Eng. Des. Sel. 25, 485, (2012)

35 Quintero-Hernández, V. et al. The change of the scFv into the Fab format improves the stability and in vivo toxin neutralization capacity of recombinant antibodies. Mol. Immunol. 44, 1307, (2007)

36 Spearman, M. et al. in Antibody Expression and Production 251 (Springer Netherlands, 2011).

37 Zheng, K. et al. The impact of glycosylation on monoclonal antibody conformation and stability. MAbs 3, 568, (2011)

38 Powers, E. T. et al. Diversity in the origins of proteostasis networks—a driver for protein function in evolution. Nat. Rev. Mol. Cell Bio. 14, 237, (2013)

39 Vieira Gomes, A. M., et al. Comparison of yeasts as hosts for recombinant protein production. Microorganisms 6, 38, (2018)

40 Matreyek, K. A. et al. Multiplex assessment of protein variant abundance by massively parallel sequencing. Nat. Genet. 50, 874, (2018)

41 Javanmardi, K. et al. Rapid characterization of spike variants via mammalian cell surface display. Mol. Cell 81, 5099, (2021)

42 Matreyek, K. A. et al. An improved platform for functional assessment of large protein libraries in mammalian cells. Nucleic Acids Res. 48, e1, (2020)

43 Tedman, A. et al. Deep receptor scanning reveals general sequence constraints on GPCR biosynthesis. bioRxiv, 2025.09.19.677468, (2025)

44 Nutalai, R. et al. Potent cross-reactive antibodies following Omicron breakthrough in vaccinees. Cell 185, 2116, (2022)

45 Tuekprakhon, A. et al. Antibody escape of SARS-CoV-2 Omicron BA. 4 and BA. 5 from vaccine and BA. 1 serum. Cell 185, 2422, (2022)

46 Kamath, N. D. et al. Multiplex functional characterization of protein variant libraries in mammalian cells with single-copy genomic integration and high-throughput DNA sequencing. Methods Mol. Biol. 2774, 135, (2024)

47 Peterman, N. et al. Sort-seq under the hood: implications of design choices on large-scale characterization of sequence-function relations. BMC Genomics 17, 1, (2016)

48 Hadfield, J. et al. Nextstrain: real-time tracking of pathogen evolution. Bioinformatics 34, 4121, (2018)

49 Abdi, H. et al. Principal component analysis. WIREs Comput. Stat. 2, 433, (2010)

50 McCandlish, D. M. Visualizing fitness landscapes. Evolution 65, 1544, (2011)

51 Kimura, M. On the probability of fixation of mutant genes in a population. Genetics 47, 713, (1962)

52 Amitai, A. et al. A population dynamics model for clonal diversity in a germinal center. Front. Microbiol. 8, 1693, (2017)

53 Mesin, L. et al. Germinal center B cell dynamics. Immunity 45, 471, (2016)

54 Morán-Tovar, R., et al. Non-equilibrium antigen recognition in acute infections. bioRxiv, doi: 10.1101/2023.04.05.535743, (2023)

55 Phillips, A. M. et al. Binding affinity landscapes constrain the evolution of broadly neutralizing anti-influenza antibodies. eLife 10, e71393, (2021)

56 Phillips, A. M. et al. Hierarchical sequence-affinity landscapes shape the evolution of breadth in an anti-influenza receptor binding site antibody. eLife 12, e83628, (2023)

57 Poelwijk, F. J. et al. The Context-Dependence of Mutations: A Linkage of Formalisms. PLOS Comput. Biol. 12, e1004771, (2016)

58 Chou, H.-H. et al. Diminishing returns epistasis among beneficial mutations decelerates adaptation. Science 332, 1190, (2011)

59 Kelly, R. L. et al. Chaperone proteins as single component reagents to assess antibody nonspecificity. MAbs 9, 1036, (2017)

60 Yang, L. et al. Antigen presentation dynamics shape the antibody response to variants like SARS-CoV-2 Omicron after multiple vaccinations with the original strain. Cell Rep. 42, 112256, (2023)

61 Forsyth, C. M. et al. Deep mutational scanning of an antibody against epidermal growth factor receptor using mammalian cell display and massively parallel pyrosequencing. MAbs 5, 523, (2013)

62 Pappas, L. et al. Rapid development of broadly influenza neutralizing antibodies through redundant mutations. Nature 516, 418, (2014)

63 Chun, J., et al. Anti-malaria antibody engineering broadens recognition motifs and reveals new homotypic interactions that enhance protective breadth. bioRxiv, (2025)

64 Kirby, M. B., et al. Retrospective SARS-CoV-2 human antibody development trajectories are largely sparse and permissive. Proc. Nat. Acad. Sci. USA 122, e2412787122, (2025)

65 Adams, R. M. et al. Epistasis in a fitness landscape defined by antibody-antigen binding free energy. Cell Syst. 8, 86, (2019)

66 Schmidt, A. G., et al. Preconfiguration of the antigen-binding site during affinity maturation of a broadly neutralizing influenza virus antibody. Proc. Nat. Acad. Sci. USA 110, 264, (2013)

67 Miton, C. M. et al. How mutational epistasis impairs predictability in protein evolution and design. Protein Sci. 25, 1260, (2016)

68 Boder, E. T. et al. Yeast surface display for directed evolution of protein expression, affinity, and stability. Methods Enzymol. 328, 430, (2000)

69 Guest, J. D. et al. An expanded benchmark for antibody-antigen docking and affinity prediction reveals insights into antibody recognition determinants. Structure 29, 606, (2021)

70 Harvey, E. P., et al. An in silico method to assess antibody fragment polyreactivity. Nat. Comm. 13, 7554, (2022)

71 Paul, S. et al. Machine learning enables efficient and effective affinity maturation of nanobodies. bioRxiv, 2026.01.11.698911, (2026)

72 Jagota, M. et al. Learning antibody sequence constraints from allelic inclusion. Cell Syst. 16, (2025)

73 Hie, B. L. et al. Efficient evolution of human antibodies from general protein language models. Nat. Biotechnol. 42, 275, (2024)

74 Tran, V. Q. et al. Rapid directed evolution guided by protein language models and epistatic interactions. Science 0, eaea1820

75 Yin, R. et al. Evaluation of AlphaFold antibody–antigen modeling with implications for improving predictive accuracy. Prot. Sci. 33, e4865, (2024)

76 Gao, M., et al. Improved deep learning prediction of antigen–antibody interactions. Proc. Nat. Acad. Sci. USA 121, e2410529121, (2024)

77 Ruffolo, J. A., et al. Fast, accurate antibody structure prediction from deep learning on massive set of natural antibodies. Nat. Comm. 14, 2389, (2023)

78 Evans, R., et al. Protein complex prediction with AlphaFold-Multimer. bioRxiv, 2021.10.04.463034, (2022)

79 Foote, J. et al. Conformational isomerism and the diversity of antibodies. Proc Natl Acad Sci USA 91, 10370, (1994). PMC45021.

80 Berman, H. M. et al. The Protein Data Bank. Nucleic Acids Res. 28, 235, (2000)

81 Li, T. et al. Rigidity Emerges during Antibody Evolution in Three Distinct Antibody Systems: Evidence from QSFR Analysis of Fab Fragments. PLOS Comput. Biol. 11, e1004327, (2015)

82 Fernández-Quintero, M. L. et al. Local and global rigidification upon antibody affinity maturation. Front. Mol. Biosci. 7, 182, (2020)

83 Ovchinnikov, V. et al. Role of framework mutations and antibody flexibility in the evolution of broadly neutralizing antibodies. eLife 7, e33038, (2018)

84 Wang, W. et al. Conformational Selection and Induced Fit in Specific Antibody and Antigen Recognition: SPE7 as a Case Study. J. Phys. Chem. B 117, 4912, (2013)

85 Ye, J. et al. IgBLAST: an immunoglobulin variable domain sequence analysis tool. Nucleic Acids Res. 41, W34, (2013)

86 Engler, C. et al. A one pot, one step, precision cloning method with high throughput capability. PLoS One 3, e3647, (2008)

87 Dadonaite, B. et al. Spike deep mutational scanning helps predict success of SARS-CoV-2 clades. Nature 631, 617, (2024)

88 Dupic, T., et al. Protein sequence landscapes are not so simple: on reference-free versus reference-based inference. bioRxiv, 2024.01.29.577800, (2024)

89 Wells, J. A. Additivity of mutational effects in proteins. Biochemistry 29, 8509, (1990)

90 Sailer, Z. R. et al. Detecting High-Order Epistasis in Nonlinear Genotype-Phenotype Maps. Genetics 205, 1079, (2017)

91 Goddard, T. D. et al. UCSF ChimeraX: Meeting modern challenges in visualization and analysis. Protein Sci. 27, 14, (2018)

92 The PyMOL Molecular Graphics System (2015).

93 Hagberg, A. et al. Exploring network structure, dynamics, and function using NetworkX. (Los Alamos National Laboratory (LANL), 2007).

94 Yen, J. Y. Finding the k shortest loopless paths in a network. Manag. Sci. 17, 712, (1971)

95 Hsieh, C.-L. et al. Structure-based design of prefusion-stabilized SARS-CoV-2 spikes. Science 369, 1501, (2020)

96 Ereño-Orbea, J. et al. Structural basis of enhanced crystallizability induced by a molecular chaperone for antibody antigen-binding fragments. J. Mol. Biol. 430, 322, (2018)

97 Punjani, A. et al. cryoSPARC: algorithms for rapid unsupervised cryo-EM structure determination. Nat. Methods 14, 290, (2017)

98 Jumper, J. et al. Highly accurate protein structure prediction with AlphaFold. Nature 596, 583, (2021)

99 Afonine, P. V. et al. Real-space refinement in PHENIX for cryo-EM and crystallography. Acta Crystallogr. D Biol. Crystallogr. 74, 531, (2018)

100 Emsley, P. et al. Features and development of Coot. Acta Crystallogr D Biol Crystallogr 66, 486, (2010)

101 Chen, V. B. et al. MolProbity: all-atom structure validation for macromolecular crystallography. Acta Crystallogr. D Biol. Crystallogr. 66, 12, (2010)

