## Supplementary material for "Biophysical trade-offs in antibody evolution are resolved by conformation-mediated epistasis": Supplemental File 5.pdf

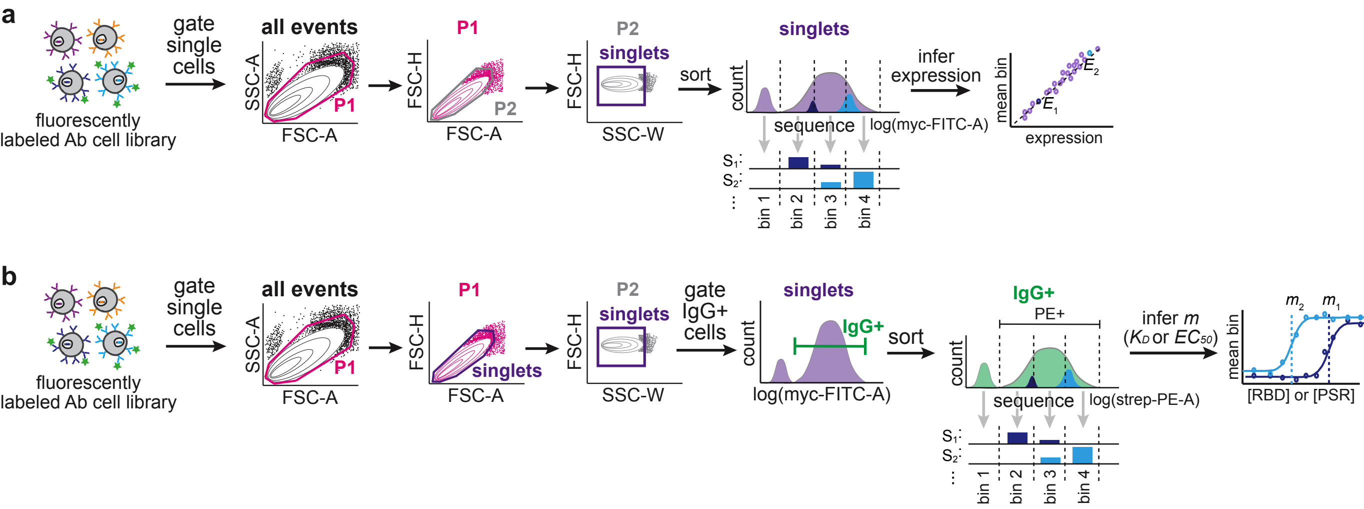


**Sorting scheme for measurements of expression (a) and binding affinity and polyspecificity (b).**
**a**, For expression, single cells were sorted into four, six, or eight populations based on FITC fluorescence intensity, with each gate capturing 25%, 16.7%, or 12.5% of the library, respectively.
**b**, For binding and polyspecificity, single IgG+ cells were sorted based on PE fluorescence using four gates – one containing non-binders and the other three containing 33% of the remaining population (low, medium, and high PE intensity).
