## Supplementary material for "Biophysical trade-offs in antibody evolution are resolved by conformation-mediated epistasis": Supplemental File 12.docx

| **Supplemental Table 1. Grid and Glacios Imaging Conditions** | | | |
| --- | --- | --- | --- |
| **Sample** | **BA.1+germline+LCKappa** | **BA.4+germline+LCKappa** | **BA.4+Omi32+LCKappa** |
| Final Sample concentrations | 4 μM BA.1 (monomer), 8 μM germline, 16 μM LCKappa | 4 μM BA.4 (monomer), 20 μM germline, 40 μM LCKappa | 4 μM BA.1, 6 μM OmiGL, 12 μM LCKappa |
| Grid | UltrAuFoil 1/1 | UltrAuFoil 0.6/1 | UltrAuFoil 0.6/1 |
| Camera | Thermofisher Falcon 4 | Thermofisher Falcon 4 | Thermofisher Falcon 4 |
| Imaging Mode | Nanoprobe | Nanoprobe | Nanoprobe |
| Detector Mode | Counting | Counting | Counting |
| Electron Voltage | 200 kV | 200 kV | 200 kV |
| Nominal defocus | -1.0 μm to -2.5 μm | -1.0 μm to -2.5 μm | -1.0 μm to -2.5 μm |
| Nominal magnification | 150,000 | 150,000 | 150,000 |
| Pixel size | 0.923 | 0.936 | 0.936 |
| Total dose | 24.85 e^-^/Å^2^ | 25.05 e^-^/Å^2^ | 25.05 e^-^/Å^2^ |
| EER internal frames | 889 | 1491 | 1491 |
| Imaging | Image shift | Image shift | Image shift |
| Total # of images | 10,318 | 5,151 | 5,466 |

| **Supplemental Table 2. Grid and Krios Imaging Conditions** | | |
| --- | --- | --- |
| **Sample** | **germline+LCKappa** | **Omi32+LCKappa** |
| Final Sample concentration | 25 μM germline, 50 μM LCKappa | 25 μM Omi32, 50 μM LCKappa |
| Grid | UltrAuFoil 0.6/1 | UltrAuFoil 0.6/1 |
| Camera | Gatan K3 | Gatan K3 |
| Imaging Mode | Nanoprobe | Nanoprobe |
| Detector Mode | Counting | Counting |
| Electron Voltage | 300 kV | 300 kV |
| Nominal defocus | -0.5 μm to -2.0 μm | -0.5 μm to -2.0 μm |
| Nominal magnification | 165,000 | 165,000 |
| Pixel size | 0.512 | 0.512 |
| Total dose | 50.13 e^-^/Å^2^ | 50.54 e^-^/Å^2^ |
| Imaging | Image shift | Image shift |
| Total # of images | 8,731 | 8,653 |

| **Supplemental Table 3. Refinement Statistics** | | | | | | | | |
| --- | --- | --- | --- | --- | --- | --- | --- | --- |
| **Sample** | **BA.1+germline+LCKappa** | | **BA.4+germline+LCKappa** | | | **BA.4+Omi32+LCKappa** | | |
| **PDB ID** | **11OQ** | | **11OO** | | | **11OL** | | |
| **EMDB ID** | **EMD-75891** | | **EMD-75889** | | | **EMD-75887** | | |
| Total extracted picks | 2,254,193 | | 1,811,974 | | | 2,320,037 | | |
| Final particles (no.) | 47,421 (C3 symmetry expanded) | | 160,551 | | | 106,182 | | |
| Symmetry | C1 | | C1 | | | C1 | | |
| FSC 0.143 | 3.2 Å | | 3.4 Å | | | 3.2 Å | | |
| **Model** | | | | | | | | |
| Chains | 4 | | 4 | | | 4 | | |
| Atoms | 5673 (Hydrogens: 0) | | 5781 (Hydrogens: 0) | | | 5801 (Hydrogens: 0) | | |
| Residues | Protein: 735 Nucleotide: 0 | | Protein: 746 Nucleotide: 0 | | | Protein: 750 Nucleotide: 0 | | |
| Water | 0 | | 0 | | | 0 | | |
| Ligands | 0 | | NAG: 1 | | | NAG: 1 | | |
| **Bonds (RMSD)** | | | | | | | | |
| Length (Å) (# > 4sigma) | 0.003 (0) | | 0.004 (0) | | | 0.005 (1) | | |
| Angles (°)(# > 4sigma) | 0.656 (1) | | 1.012 (0) | | | 1.024 (3) | | |
| MolProbity score | 1.74 | | 1.68 | | | 1.59 | | |
| Clash score | 8.36 | | 5.91 | | | 5.37 | | |
| **Ramachandran plot (%)** | | | | | | | | |
| Outliers | 0.14 | | 0.14 | | | 0.13 | | |
| Allowed | 3.99 | | 5.03 | | | 4.18 | | |
| Favored | 95.87 | | 94.84 | | | 95.69 | | |
| **Ramachandran plot Z-score (RMSD)** | | | | | | | | |
| Whole | -0.54 (0.32) | | -0.64 (0.32) | | | -0.31 (0.32) | | |
| Helix | -3.26 (0.58) | | -3.06 (0.61) | | | -1.81 (0.96) | | |
| Sheet | 0.06 (0.32) | | -0.32 (0.30) | | | -0.16 (0.33) | | |
| Loop | -0.17 (0.32) | | 0.02 (0.34) | | | 0.00 (0.31) | | |
| Rotamer outliers (%) | 0.00 | | 0.47 | | | 0.63 | | |
| Cbeta outliers (%) | NA | | NA | | | NA | | |
| **Peptide plane (%)** | | | | | | | | |
| Cis proline/general | 13.2/0.0 | | 15.4/0.0 | | | 12.8/0.0 | | |
| Twisted proline/general | 0.0/0.0 | | 0.0/0.0 | | | 0.0/0.0 | | |
| CaBLAM outliers (%) | 2.36 | | 2.75 | | | 2.18 | | |
| **ADP (B-factors)** | | | | | | | | |
| Iso/Aniso (#) | 5673/0 | | 5781/0 | | | 5801/0 | | |
| Min/max/mean |  | |  | | |  | | |
| Protein | 17.99/149.43/64.67 | | 19.69/178.47/75.13 | | | 12.70/172.38/73.73 | | |
| Nucleotide | --- | | --- | | | --- | | |
| Ligand | --- | | 112.59/142.42/134.31 | | | 68.63/104.95/95.22 | | |
| Water | --- | | --- | | | --- | | |
| **Occupancy** | | | | | | | | |
| Mean | 1.00 | | 1.00 | | | 1.00 | | |
| occ = 1 (%) | 99.89 | | 99.57 | | | 99.52 | | |
| 0 < occ < 1 (%) | 0.11 | | 0.38 | | | 0.48 | | |
| occ > 1 (%) | 0.00 | | 0.00 | | | 0.00 | | |
| **Data** |  | |  | | |  | | |
| Box |  | |  | | |  | | |
| Lengths (Å) | 81.97, 84.73, 115.12 | | 76.75, 77.69, 124.49 | | | 75.82, 83.30, 123.55 | | |
| Angles (°) | 90.00, 90.00, 90.00 | | 90.00, 90.00, 90.00 | | | 90.00, 90.00, 90.00 | | |
| Supplied resolution (Å) | 3.2 | | 3.3 | | | 3.2 | | |
| Resolution estimates (Å) | Masked | Unmasked | | Masked | Unmasked | | Masked | Unmasked |
| d FSC (half maps; 0.143) | 3.2 | 3.4 | | 3.4 | 3.5 | | 3.3 | 3.4 |
| d 99 (full/half1/half2) | 3.3/1.9/1.9 | 3.2/1.9/1.9 | | 3.6/1.9/1.9 | 3.5/1.9/1.9 | | 3.4/1.9/1.9 | 3.3/1.9/1.9 |
| d model | 3.3 | 3.3 | | 3.6 | 3.6 | | 3.5 | 3.5 |
| d FSC model (0/0.143/0.5) | 3.0/3.1/3.3 | 3.1/3.2/3.5 | | 3.2/3.3/3.6 | 3.3/3.4/3.7 | | 3.0/3.1/3.4 | 3.1/3.2/3.5 |
| Map min/max/mean | -0.29/0.44/0.01 | | -0.40/0.72/0.00 | | | -0.35/0.54/0.00 | | |
| **Model vs. Data** | | | | | | | | |
| CC (mask) | 0.81 | | 0.76 | | | 0.78 | | |
| CC (box) | 0.62 | | 0.62 | | | 0.59 | | |
| CC (peaks) | 0.56 | | 0.55 | | | 0.52 | | |
| CC (volume) | 0.79 | | 0.74 | | | 0.76 | | |
| Mean CC for ligands | --- | | 0.34 | | | 0.62 | | |

| **Supplemental Table 4. Refinement Statistics** | | | | | |
| --- | --- | --- | --- | --- | --- |
| **Sample** | **germline+LCKappa** | | **Omi32+LCKappa** | | |
| **PDB ID** | **11OU** | | **11OR** | | |
| **EMDB ID** | **EMD-75893** | | **EMD-75892** | | |
| Total extracted picks | 2,740,105 | | 2,224,101 | | |
| Final particles (no.) | 73,477 | | 71,518 | | |
| Symmetry | C1 | | C1 | | |
| FSC 0.143 | 3.1 Å | | 3.2 Å | | |
| **Model** |  | |  | | |
| Chains | 3 | | 3 | | |
| Atoms | 4166 (Hydrogens: 0) | | 4128 (Hydrogens: 0) | | |
| Residues | Protein: 549 Nucleotide: 0 | | Protein: 545 Nucleotide: 0 | | |
| Water | 0 | | 0 | | |
| Ligands | 0 | | 0 | | |
| **Bonds (RMSD)** | | | | | |
| Length (Å) (# > 4sigma) | 0.002 (0) | | 0.003 (0) | | |
| Angles (°)(# > 4sigma) | 0.471 (0) | | 0.551 (0) | | |
| MolProbity score | 1.37 | | 1.50 | | |
| Clash score | 4.90 | | 7.54 | | |
| **Ramachandran plot (%)** | | | | | |
| Outliers | 0.00 | | 0.00 | | |
| Allowed | 2.59 | | 2.42 | | |
| Favored | 97.41 | | 97.58 | | |
| **Ramachandran plot Z-score (RMSD)** | | | | | |
| Whole | 0.48 (0.39) | | 0.31 (0.39) | | |
| Helix | 1.94 (1.37) | | 0.29 (1.13) | | |
| Sheet | 0.59 (0.35) | | 0.03 (0.34) | | |
| Loop | 0.09 (0.41) | | 0.60 (0.43) | | |
| Rotamer outliers (%) | 0.00 | | 0.89 | | |
| Cbeta outliers (%) | NA | | NA | | |
| **Peptide plane (%)** | | | | | |
| Cis proline/general | 17.9/0.0 | | 17.9/0.0 | | |
| Twisted proline/general | 0.0/0.0 | | 0.0/0.0 | | |
| CaBLAM outliers (%) | 1.69 | | 1.51 | | |
| **ADP (B-factors)** | | | | | |
| Iso/Aniso (#) | 4166/0 | | 4128/0 | | |
| Min/max/mean |  | |  | | |
| Protein | 11.91/116.02/55.48 | | 6.66/109.94/50.90 | | |
| Nucleotide | --- | | --- | | |
| Ligand | --- | | --- | | |
| Water | --- | | --- | | |
| **Occupancy** | | | | | |
| Mean | 1.00 | | 1.00 | | |
| occ = 1 (%) | 99.93 | | 99.85 | | |
| 0 < occ < 1 (%) | 0.00 | | 0.15 | | |
| occ > 1 (%) | 0.00 | | 0.00 | | |
| **Data** |  | |  | | |
| Box |  | |  | | |
| Lengths (Å) | 56.17, 76.41, 96.65 | | 56.17, 75.90, 94.62 | | |
| Angles (°) | 90.00, 90.00, 90.00 | | 90.00, 90.00, 90.00 | | |
| Supplied resolution (Å) | 3.2 | | 3.2 | | |
| Resolution estimates (Å) | Masked | Unmasked | | Masked | Unmasked |
| d FSC (half maps; 0.143) | 3.2 | 3.3 | | 3.2 | 3.3 |
| d 99 (full/half1/half2) | 3.3/1.0/1.0 | 3.2/1.0/1.0 | | 3.4/1.0/1.0 | 3.3/1.0/1.0 |
| d model | 3.4 | 3.4 | | 3.4 | 3.4 |
| d FSC model (0/0.143/0.5) | 3.0/3.1/3.3 | 3.0/3.1/3.4 | | 3.0/3.1/3.3 | 3.1/3.2/3.4 |
| Map min/max/mean | -0.32/0.40/0.00 | | -0.25/0.41/0.01 | | |
| **Model vs. Data** | | | | | |
| CC (mask) | 0.83 | | 0.83 | | |
| CC (box) | 0.66 | | 0.70 | | |
| CC (peaks) | 0.65 | | 0.68 | | |
| CC (volume) | 0.80 | | 0.80 | | |
| Mean CC for ligands | --- | | --- | | |
