## Supplementary material for "Biophysical trade-offs in antibody evolution are resolved by conformation-mediated epistasis": Supplemental_File_13_Legend.docx

**Supplemental Video 1. Antibody preconfiguration and antigen binding**. The HCDR2 and LCDR1 loops in the germline antibody (blue) undergo a conformational change upon acquiring the mutations present in Omi32 (red). Loops are shown in lighter shades of blue and red, with the HCDR2 on the left and the LCDR1 on the right. After preconfiguration, Omi32 binds RBD (grey; Omi32+BA4 is shown).
